# A Human Liver Cell Atlas: Revealing Cell Type Heterogeneity and Adult Liver Progenitors by Single-Cell RNA-sequencing

**DOI:** 10.1101/649194

**Authors:** Nadim Aizarani, Antonio Saviano, Sagar, Laurent Mailly, Sarah Durand, Patrick Pessaux, Thomas F. Baumert, Dominic Grün

## Abstract

The human liver is an essential multifunctional organ, and liver diseases are rising with limited treatment options. However, the cellular complexity and heterogeneity of the liver remain poorly understood. Here, we performed single-cell RNA-sequencing of ~5,000 cells from normal liver tissue of 6 human donors to construct the first human liver cell atlas. Our analysis revealed previously unknown sub-types among endothelial cells, Kupffer cells, and hepatocytes with transcriptome-wide zonation of these populations. We show that the EPCAM^+^ population is highly heterogeneous and consists of hepatocyte progenitors, cholangiocytes and a *MUC6*^+^ stem cell population with a specific potential to form liver organoids. As proof-of-principle, we applied our atlas to unravel phenotypic changes in cells from hepatocellular carcinoma tissue and to investigate cellular phenotypes of human hepatocytes and liver endothelial cells engrafted into a humanized *FAH*^-/-^ mouse liver. Our human liver cell atlas provides a powerful and innovative resource enabling the discovery of previously unknown cell types in the normal and diseased liver.

## MAIN

The liver is a key organ in the human body. It serves as a central metabolic coordinator with a wide array of essential functions, including regulation of glucose and lipid metabolism, protein synthesis, bile synthesis and secretion, as well as biotransformation of metabolites and xenobiotics. Furthermore, the liver is a visceral organ that is capable of remarkable natural regeneration after tissue loss^1^. However, prevalence and mortality of liver disease have risen dramatically within the last decades, and hepatocellular carcinoma (HCC) has become the second leading cause of cancer-associated death^2,3^. A key reason for the absence of efficient therapies for a number of liver diseases is the complexity of the organ and its multiple under-characterized cell types contributing to disease pathogenesis. The cellular landscape of the liver has barely been explored at single-cell resolution, which limits our molecular understanding of liver function and disease biology. A major challenge has been the difficulty to isolate viable single hepatocytes from patient tissues.

In recent years, a number of advanced single-cell methods has been developed to sequence the transcriptomes of thousands of individual cells with high sensitivity^4^. The transcriptome of a cell is highly informative on its state and can be used as the basis for the identification of cell types in health and disease^5-8^.

To characterize human liver cell types at single-cell resolution, we developed a robust pipeline for single-cell RNA-sequencing (scRNA-seq) of cryopreserved and freshly isolated patient-derived human liver samples and assembled the first human liver cell atlas consisting of 5,416 cells from six donors. We performed in-depth analysis of all liver cell types with a major focus on the stem cell compartment.

As proof-of-principle experiments we demonstrate the value of our atlas by revealing changes in cellular phenotypes in hepatocellular carcinoma (HCC) tissue and of human liver endothelial cells and hepatocytes transplanted into a humanized FAH^-/-^ mouse model.

### Establishing scRNA-seq of human liver cells to assemble a cell type atlas of the adult human liver

We applied mCEL-Seq2^9,10^ for scRNA-seq of cells from non-diseased liver tissue obtained from six patients who underwent liver resections for colorectal cancer metastasis or cholangiocarcinoma without history of chronic liver disease (Fig. 1a, Methods). We sorted and sequenced viable cells in an unbiased fashion as well as specific cell populations on the basis of cell surface markers (Extended Data Fig. 1, Methods). Since fresh liver tissue material is scarce and difficult to preserve, and biobanks with cryopreserved liver tissue derived from patients represent rich resources, we attempted to generate single-cell transcriptome data from cryopreserved cells in addition to single cell suspensions from freshly prepared liver resection specimens (Methods). We then used RaceID3 for the identification of cell types (Methods)^5^.

**Figure 1.**
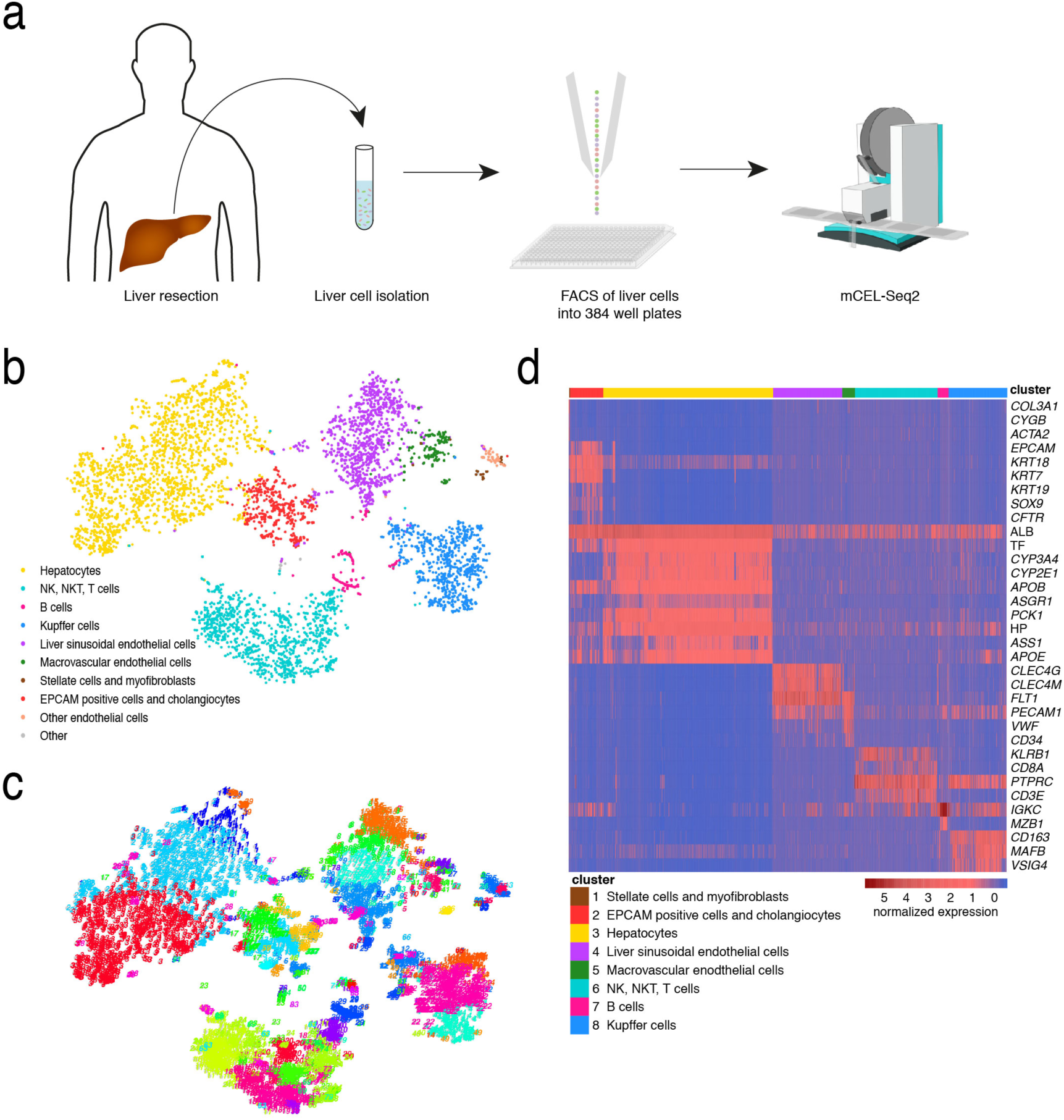
ScRNA-seq reveals cell types in the adult human liver. **a**, An outline of the protocol used for scRNA-seq of human liver cells. Specimens from liver resections were first digested to prepare single cell suspensions. Cells were stained with a viability dye and antibodies and then sorted by FACS into 384 well plates, which were later processed according to the mCEL-Seq2 protocol. **b**, t-SNE map of single-cell transcriptomes from normal liver tissue of six different donors showing the main liver cell compartments. **c**, t-SNE map of single-cell transcriptomes highlighting the clusters generated by RaceID3, revealing sub-type heterogeneity in all major cell populations of the human liver. **d**, Heatmap showing the expression of established marker genes for each cell compartment. Scale bar shows log_2_ transformed z-scores with a threshold cutoff of 6.

Cells from different patients, isolated from freshly prepared (P301, P304) or cryopreserved (P301, P304, P308, P309, P310, P311) single-cell suspensions generally co-clustered (Extended Data Fig. 1), showing that our pipeline is robust and can be used for sequencing cells from cryopreserved samples.

Based on the expression of marker genes, our atlas reveals all the main liver cell types, including hepatocytes, EPCAM^+^ bile duct epithelial cells (cholangiocytes) and liver stem cells, CLEC4G^+^ liver sinusoidal endothelial cells (LSECs), CD34^+^PECAM^high^ macro-vascular endothelial cells (MaVECs), hepatic stellate cells and myofibroblasts, Kupffer cells, NKT cells, NK cells, T cells and B cells (Fig. 1b-d). Several clusters corresponding to distinct sub-types were identified within these major populations (Fig. 1c), indicating previously unknown heterogeneity across liver cell types.

### Heterogeneity and zonation of endothelial cells, Kupffer cells, and hepatocytes

Hepatocytes are spatially heterogeneous and sub-specialized, or zonated, along the portal-central axis of the liver lobule^11-15^. Based on the metabolic sub-specialization of hepatocytes, the liver lobule has been divided into three zones; the periportal zone nearest to the portal vein, hepatic artery and bile duct, the central zone nearest to the central vein, and the mid zone located between the central and periportal zones ^11,13,15^. While previous observations had hinted at underlying zonation and sub-specialization of non-parenchymal cells like LSECs and Kupffer cells^11,16,17^, spatial heterogeneity of these cell types has remained elusive, and most studies were carried out in rodents. In order to characterize the heterogeneity of these cell types in human, we first performed differential gene expression analysis between the clusters within the endothelial and Kupffer cell compartments, respectively (Methods).

Our analysis revealed two major populations of endothelial cells in the human liver, the MaVECs and LSECS. LSECs line the sinusoids of the liver lobule and are *CLEC4G*^+^*PECAM1*^low^, while MaVECs line the hepatic arteries and veins and are *CD34*^+^*PECAM1*^high^ cells^18,19^ (Extended Data Fig. 2). LSECs have different functions and morphology from venous and arterial endothelial cells, i.e. they are fenestrated and have filtering and scavenging roles^17,20-22^. However, gene expression differences between MaVECs and LSECs are only known for few markers, such as CD34 and PECAM1^19^. Differential gene expression analysis between *CD34*^+^ MaVECs (clusters 5 and 41) and *CLEC4G*^+^ LSECs (clusters 7, 8, 9, 15) revealed novel markers for these two types, e.g. the water channel *AQP1* for *CD34*^+^ endothelial cells and *FCN3* for LSECs (Extended Data Fig. 2a,c). Within the *CD34*^+^*AQP1*^+^ compartment we identified a *CPE*^+^ (cluster 41) and a *CPE*^-^ (cluster 5) sub-population (Extended Data Fig. 2d). In addition, we identified several novel sub-types, including a *CCL21*^+^ population, which expresses angiogenesis-associated genes like the netrin receptor *UNC5B*^23,24^ and *TFF3^25^* (cluster 25) (Extended Data Fig. 2c). Another population, highly expresses *H19*, a non-coding RNA involved in early development^26^, together with both LSEC- and MaVSEC-specific genes (cluster 26), corresponding to a potential pan-endothelial cell progenitor (Extended Data Fig. 2e).

Cluster-to-cluster analysis in the *CD163*^+^ Kupffer cell compartment revealed three main subsets: a *VCAM1*^+^*LIRB5*^+^*MARCO*^+^ subset, a *CD1C*^+^ subset, and a *VCAN*^+^ subset (Extended Data Fig. 3a).

Differential gene expression and pathway enrichment analysis reveals an M2-like anti-inflammatory gene signature for *VCAM1*^+^*LILRB5*^+^ Kupffer cells while *CD1C*^+^ Kupffer cells exhibit an M1-like pro-inflammatory gene signature with high expression of genes involved in MHC Class II antigen presentation, suggesting the presence of M1-/M2-like Kupffer cell subsets in the steady state normal liver.

ScRNA-seq has been shown to be highly informative on hepatocyte zonation in mouse liver^27^. In order to infer transcriptome-wide zonation across liver cell types, we reasoned that the major axis of variability for each cell type could reflect gene expression changes associated with zonation. Hence, we ordered LSECs, Kupffer cells, and hepatocytes by diffusion pseudo-time (dpt)^28^ along this axis (Methods) and applied self-organizing maps (SOMs) on dpt-ordered cells to infer co-expression modules along this axis (Fig. 2a). This approach recovered known zonated expression patterns of landmark genes in human hepatocytes, e.g. *ALB* and *PCK1* (module 11, highly expressed in the periportal zone), *HP* (module 4, up-regulated in the mid-zone), and *GLUL* and *CYP1A2*, (module 5, expressed in the central zone) (Fig. 2a, Extended Data Fig. 3b). Pathway enrichment analysis of the hepatocyte modules reveals that periportal hepatocytes are enriched for genes involved in biological oxidations consistent with a gradient of oxygen concentration peaking in the periportal zone ^11,13,15^(Fig. 2a, Methods). These results validate the use of dpt for inferring transcriptome-wide zonation of liver cells.

**Figure 2.**
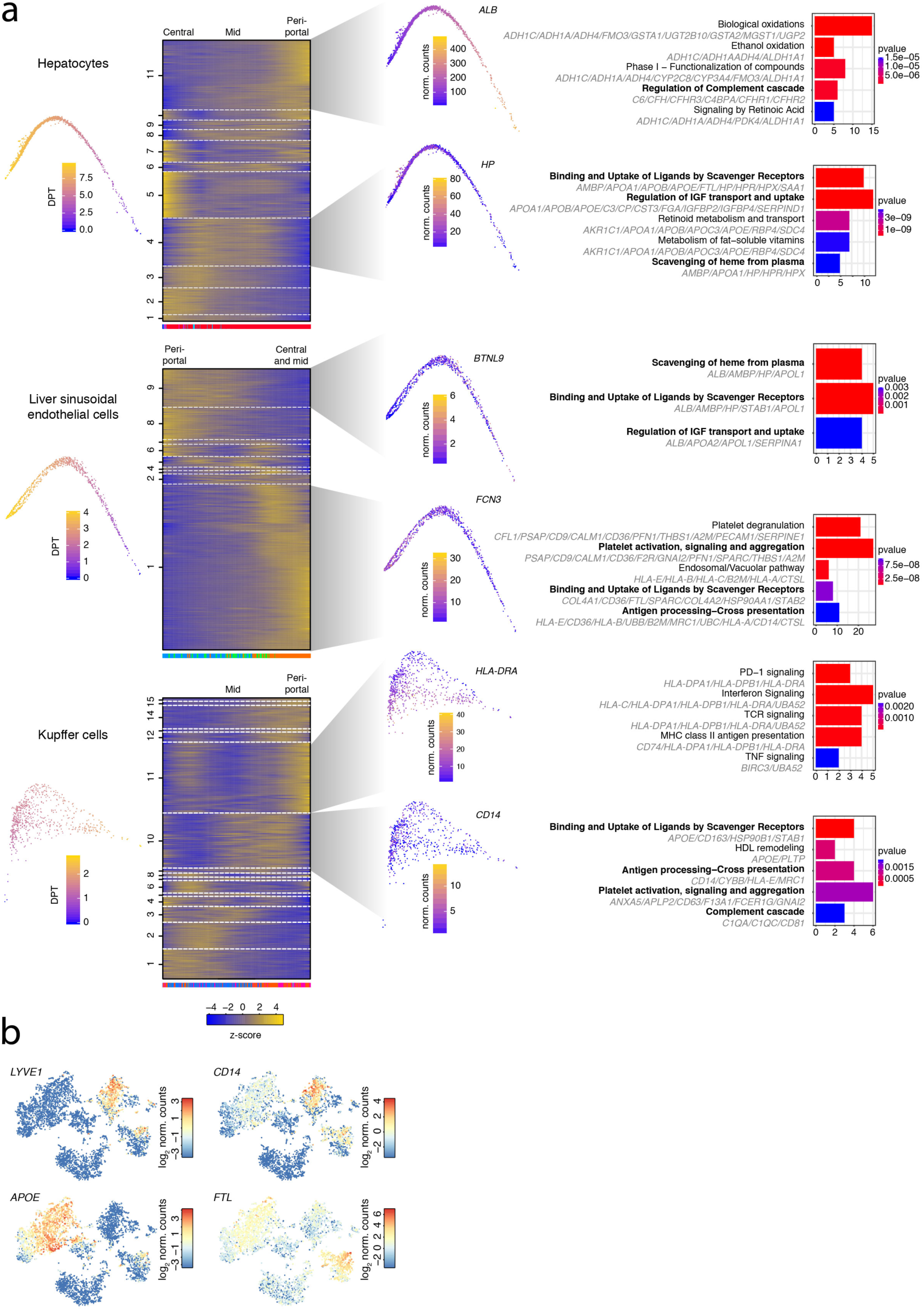
Heterogeneity and zonation of hepatocytes, LSECs and Kupffer cells. **a**, Diffusion maps (left) and self-organizing maps (SOM, middle) of the pseudo-temporally ordered single-cell transcriptomes for hepatocytes (top), LSECs (middle), and Kupffer cells (bottom), indicative of cell type zonation in the human liver lobule. Expression modules were derived as described previously^10^ (Methods). Pathways enriched for the genes in particular modules are shown to the right. Pathways highlighted in bold indicate pathways co-zonated across cell types. The color bar at the bottom of the SOMs indicates the cluster of origin for each cell. The number on the y-axes label modules of co-expressed genes. **b**, Expression t-SNE maps of the co-zonated genes *LYVE1*, *CD14*, *APOE* and *FTL* in hepatocytes, LSECs, and Kupffer cells.

LYVE1 and CD14 were recently identified as markers distinguishing two LSEC subsets, the CD14^+^LYVE1^+^ subset residing in the mid and central zones and the CD14^-^LYVE1^-^ subset residing in the periportal zone^29^. However, our analysis shows that *CD14* and *LYVE1* are co-expressed by LSEC and Kupffer cell subsets, indicating their co-zonation in the mid and central zones and the existence of co-zonated genes across these different cell types (Fig. 2b). SOMs for LSECs and Kupffer cells reveal that the transcriptomes of *CD14*^+^ mid and central zonal cells and of *CD14*^-^ periportal cells are assigned to separate modules (Fig. 2a). Mid and central zonal LSECs (module 1) highly express *FCN3* and *CTSL* in a gradient-like pattern while periportal LSECs (modules 8 and 9) highly express *BTNL9* and the angiogenesis, receptor scavenging and lymphocyte homing-associated gene *STAB1*^30^ (Fig. 2a, Extended Data Fig. 3c). *CD14*^+^ mid-zonal Kupffer cells include the *VCAM1*^+^*LILRB5*^+^*MARCO*^+^ M2-like subset, while the *CD14*^-^ Kupffer cells correspond to the *CD1C*^+^*HLA-DRA*^high^ M1-like sub-type (Fig. 2a,b, Extended Data Fig. 3a).

After inferring the zonation of the different subsets in LSECs, Kupffer cells and hepatocytes, we performed, for each of these cell types, a pathway enrichment analysis on genes from the modules that correspond to the different lobular zones. Interestingly, our analysis revealed novel pathways that are shared across all three cell types, including co-zonated pathways such as binding and uptake of ligands by scavenger receptors, which is shared across mid-zonal hepatocytes, LSECs, and Kupffer cells (Fig. 2a, Extended Data Fig. 4). As another example, antigen processing and cross presentation, is co-zonated across *CD14*^+^*LYVE1*^+^ midzonal LSECs and Kupffer cells. Particular genes within these pathways exhibit co-zonation across these cell types, e.g. *FTL*, *APOE* and *HLA-E*, *LYVE1* and *CD14* (Fig. 2b). Interestingly, we also observe potential co-zonation of ribosomal gene expression across these cell types (Extended Data Fig. 5).

Although we were not able to observe zonation of B cells, we identified the gene signatures of two B cell subsets; an MS4A1^+^CD37^+^ subset, which corresponds to circulating B cells upregulating MHC class II components, and a liver resident MZB1^+^ B cell subset (Extended Data Fig. 6).

In summary, our single cell analysis unravels transcriptome-wide zonation of human hepatocytes and non-parenchymal cells and suggests that subsets of Kupffer cells, LSECs and hepatocytes share similar functions and gene expression profiles as a result of their zonation in the liver lobule, indicating functional co-operation across cell types in the human liver.

### Identification of a putative stem cell in the EPCAM^+^ compartment of the normal adult human liver

Liver regeneration after tissue damage involves the replication of several liver cell types including hepatocytes, cholangiocytes and LSECs. Furthermore, different types of liver damage lead to specific mechanisms of liver regeneration. For example, mouse cholangiocytes have been shown to transdifferentiate into hepatocytes after impaired hepatocyte regeneration^31^, and rat hepatocytes transdifferentiate into cholangiocytes following bile duct ligation and toxic biliary injury^32^. Moreover, in mouse, a subset of cholangiocytes was identified to have clonogenic potential and to differentiate into hepatocytes^33^. However, the existence of a naïve adult stem cell population in the human liver and its contribution to turnover and regeneration remains controversial. In the adult human liver, rare EPCAM^+^ cells have been termed hepatic stem cells^34^. These cells can form dense round colonies when cultured and have been reported to be bi-potent progenitors of hepatoblasts, which differentiate into cholangiocytes or hepatocytes both *in vitro* and *in vivo*^34,35^.

In search for the genuine liver stem cell, we sorted and sequenced single EPCAM^+^ cells from adult human livers. The EPCAM^+^ cells that we sequenced were negative for proliferation markers *MKI67* and *PCNA* and the hepatoblast marker *AFP* (Extended Data Fig. 7a). This suggests that the heterogeneity in the EPCAM^+^ compartment is not arising as a result of proliferation but that the observed sub-types reside in the normal human liver.

Using our approach, we discovered novel liver stem cell surface markers whose expression correlates with that of *EPCAM* or *TROP1*; these include *TACSTD2* (*TROP2)*, *CD24*, *FGFR2*, *AQP1*, *DEFB2*, *TM4SF4* and *CLDN4* (Fig. 3a and Extended Data Fig. 7b). Flow cytometry profiles of EPCAM and the novel marker TROP2 resolve the presence of a TROP2^high^ and a TROP2^low^ population within the EPCAM^+^ compartment (Fig. 3b). Moreover, forward and side-scatter profiles of EPCAM^+^ cells indicate that the compartment is heterogeneous and consists of populations with different sizes and morphologies (Fig. 3b).

**Figure 3.**
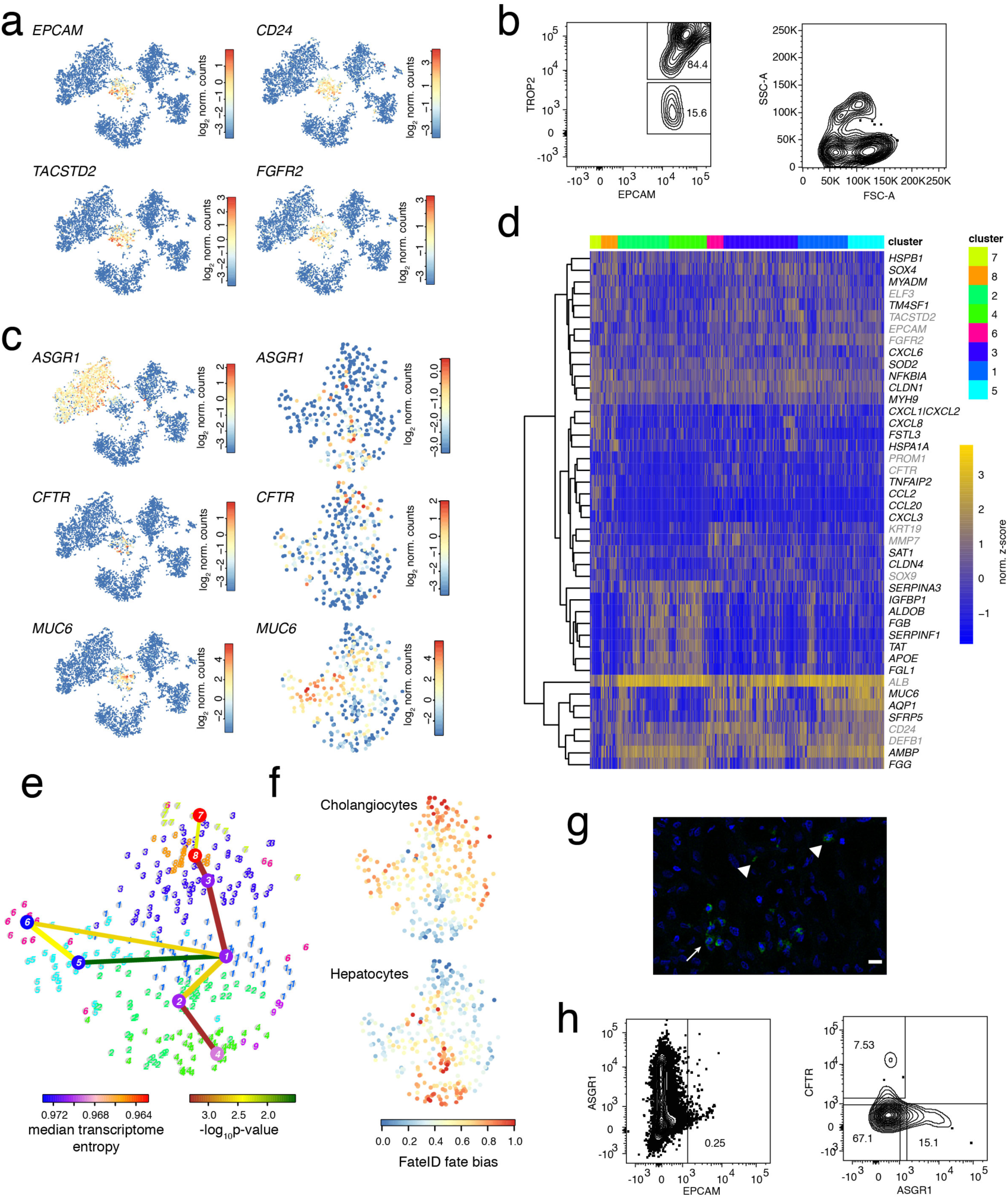
Identification of a putative stem cell population in the adult human liver. **a**, t-SNE maps showing the expression of the hepatic stem cell marker *EPCAM* and the novel markers *TACSTD2*, *CD24* and *FGFR2* whose expression correlates with that of *EPCAM*. **b**, FACS plots for sorted EPCAM^+^ cells showing EPCAM and TROP2 expression (left) and forward and side scatter (right). **c**, Expression t-SNE maps of *ASGR1*, *CFTR* and *MUC6*, revealing three major populations in the *EPCAM*^+^ positive compartment: a hepatocyte biased progenitor population, a cholangiocyte biased population, and a *MUC6*^+^ putative stem cell population. **d**, Heatmap showing differentially expressed genes between the hepatocyte progenitors (cluster 4), the cholangiocyte biased population (cluster 7), and the *MUC6*^+^ putative stem cell population (cluster 5) from a separate RaceID3 analysis of the *EPCAM*^+^ compartment. Differentially expressed genes were inferred by pairwise differential gene expression analysis between the fate-biased clusters and the *MUC6*^+^ cluster. (Benjamini-Hochberg corrected *P*<0.01). Only genes with a mean expression >2 in at least one of the clusters and a log_2_-fold changes of >2 in at least one of the comparisons were included. Additional informative marker genes are highlighted in grey. Scale bar shows log_2_ transformed z-scores. **e**, StemID2^38^ analysis of the *EPCAM*^+^ compartment reveals that hepatocyte- and cholangiocyte-biased populations share a common origin and bifurcate from the *MUC6*^+^ cells (cluster 5), which have the highest transcriptome entropy. The link color represents significance. Shown are links with *P*<0.05. The node color highlights transcriptome entropy. **f**, FateID analysis of the *EPCAM*^+^ compartment highlights populations that are preferentially biased towards hepatocyte progenitors and cholangiocytes, respectively, and reveals similar bias towards both lineages in *MUC6*^+^ cells. **g**, Immunofluorescence staining of MUC6 (green) shows localization of MUC6^+^ cells in the bile ducts (white arrow) and canals of Herring. Arrow heads indicate individual MUC6+ cells. DAPI staining is shown in blue. Scale bar, 10 μm. **h**, FACS plots of ASGR1 and CFTR expression in the EPCAM^+^ compartment.

Our single-cell transcriptome analysis revealed that the compartment consists of three main populations: (1) a hepatocyte progenitor population (cluster 17), which lowly expresses ductal markers like *EPCAM* and *TACTSD2* and hepatocyte genes like *TF* and *ASGR1* at low levels compared to mature hepatocytes, (2) a *CFTR*^+^*FST3*^+^ cholangiocyte population (cluster 48), and (3) a *MUC6*^+^ population (cluster 13) (Fig. 3c,d and Extended Data Fig. 7c). The observation of cholangiocyte- and hepatocyte-biased populations in the EPCAM^+^ compartment suggested that the remaining *MUC6*^+^ population, which also expresses the WNT signaling modulator *SFRP5*, could be a putative naïve stem cell population. Interestingly, *MUC6*, a major gastric mucin, is expressed by pancreatic progenitors stemming from the multi-potent bile duct tree stem cells^36,37^, which have been proposed to be the origin of the EPCAM^+^ hepatic stem cells, providing supporting evidence for a potential stem cell nature of the *MUC6*^+^ cells.

We then re-analyzed only the EPCAM^+^ population with RaceID3 and employed our recent StemID2 algorithm for lineage reconstruction and stem cell prediction^38^ (Fig. 3c-e, Methods). This analysis revealed that the *MUC6*^+^ population (clusters 5 and 6) maximizes the transcriptome entropy, as expected for stem cells^38^, and bifurcates into hepatocyte progenitors and cholangiocytes (Fig. 3e). To better understand the emergence of fate bias towards the two lineages, we applied the FateID algorithm^10^. Consistently, this algorithm infers similar probabilities of the *MUC6*^+^ population to differentiate towards hepatocytes and cholangiocytes, while all other clusters within the EPCAM^+^ population show a distinct bias towards a particular lineage (Fig. 3f).

We validated the existence of the *MUC6*^+^ progenitor population in the human liver tissue by antibody staining and found that these cells reside in bile duct structures and the canals of Herring (Fig. 3g), which have been suggested as candidate sites harboring hepatic stem cells^35,39^. We note that upregulation of MUC6 in proliferating bile ductular cells was observed in diseased liver^40^, particularly in chronic viral hepatitis and necro-inflammation, indicating potential activation and proliferation of *MUC6*^+^ progenitors as a result of liver damage.

In order to validate our prediction, we first confirmed the presence of the three populations by FACS on the basis of EPCAM, ASGR1 and CFTR expression (Fig. 3h), and then sorted each population for organoid culturing^41^. The only cells that grew and formed organoids were negative for the expression of the hepatocyte progenitor and cholangiocyte markers, corresponding to the *MUC6*^+^ population (Fig. 4a). After organoid culturing, we stained for EPCAM and ASGR1 and identified an EPCAM^+^ASGR1^+^ population within organoids, akin to the hepatocyte progenitor population from the patients (Fig. 4b). We also counted few cells, which have lost their EPCAM expression and likely correspond to differentiated cell types.

**Figure 4.**
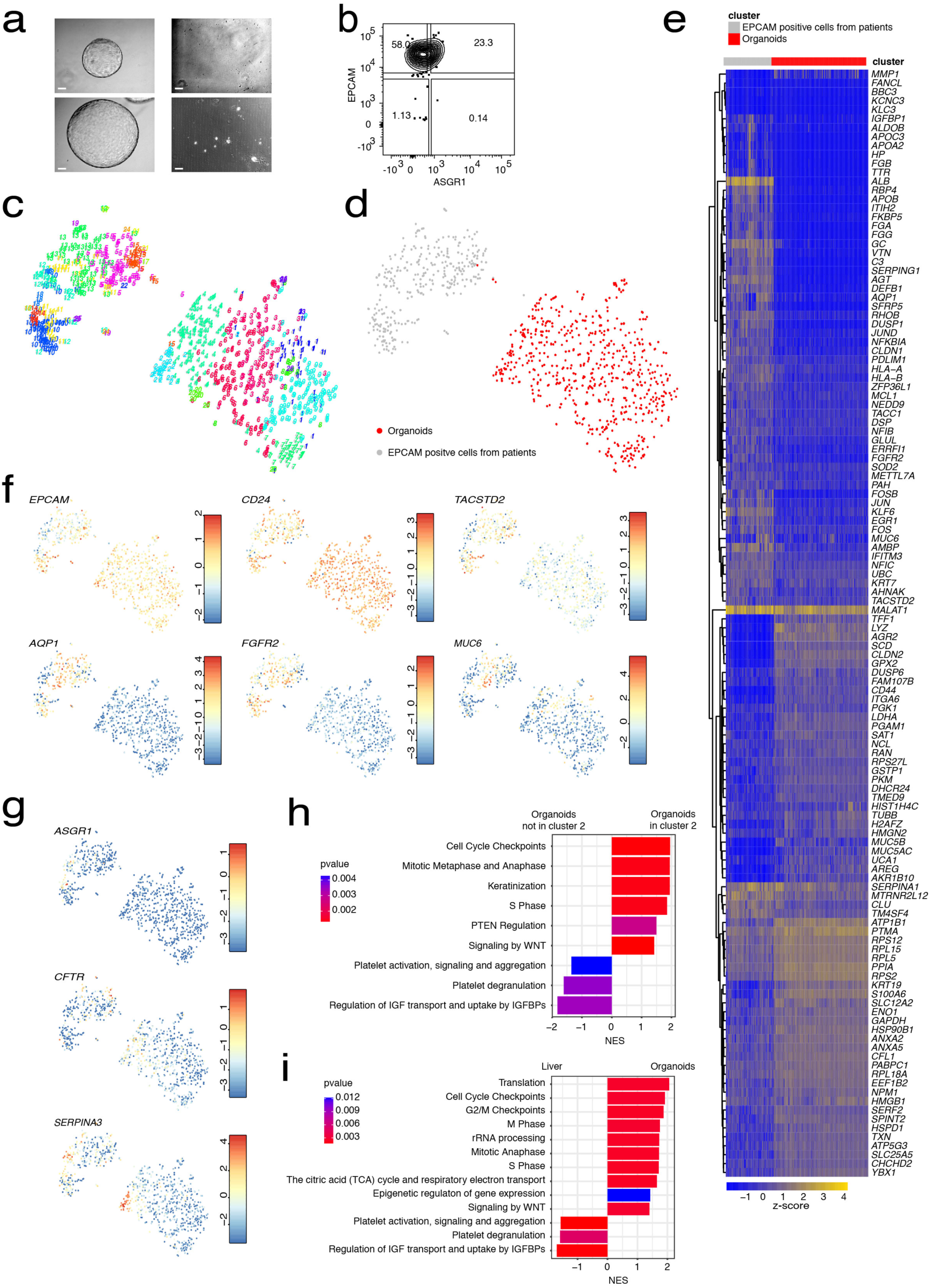
*MUC6*^+^ putative stem cells are a specific source of liver organoid formation. **a**, Organoid culturing of the three populations from the EPCAM^+^ compartment. Organoids were only formed from the EPCAM^+^ compartment negative for hepatocyte and cholangiocyte markers, corresponding to the *MUC6*^+^ population (left). ASGR1^+^ (right, upper panel) or CFTR^+^ (right, lower panel) populations did not give rise to organoids. Images were taken 13 days after sorting the cells for culture in n=2 independent experiments. Scale bars correspond to 500 μm. **b**, FACS plot of EPCAM and ASGR1 expression in cells from the organoids. **c**, t-SNE map showing RaceID3 clusters for *EPCAM*^+^ cells from the patients and the organoids. **d**, Symbol t-SNE map showing *EPCAM*^+^ cells from the patients and the organoids. **e**, Heatmap of differentially expressed genes between patient cells and organoid cells. Differentially expressed genes were inferred by pairwise differential gene expression analysis between patient cells and organoid cells (Benjamini-Hochberg corrected *P*<0.01). Genes with mean expression >1 and log_2_-fold change >1 were included. **f**, Expression t-SNE maps of the stem cell markers *EPCAM*, *CD24*, *TACSTD2*, *AQP1, FGFR2* and *MUC6*. **g**, Expression t-SNE maps of the mature cell type markers *ASGR1*, *CFTR* and *SERPINA3*. **h**, Gene set enrichment analysis (GSEA) of differentially expressed genes between human liver cells (Liver) and cells from the organoids (Organoids). Differentially expressed genes were inferred by pairwise differential gene expression analysis between patient cells and organoid cells (Benjamini-Hochberg corrected *P*<0.01). **i**, GSEA of differentially expressed genes between cells from cluster 2 (Organoids cluster 2) and the remaining clusters from the organoids (Organoids not cluster 2). Differentially expressed genes were inferred by pairwise comparison between cluster 2 and other organoid clusters (Benjamini-Hochberg corrected *P*<0.01). The bar charts in (h) and (i) show the normalized enrichment score (NES) and highlight the p-value.

To elucidate the cell type composition of the organoids in depth, we performed scRNA-seq. Co-analysis of the organoid cells and the EPCAM^+^ cells sequenced directly from the patients demonstrates marked transcriptome differences (Fig. 4c,d). While *EPCAM* and *CD24* were expressed in cells from the organoids and patients, organoid cells downregulated various markers expressed in the patients’ EPCAM^+^ cells such as *AQP1* and *SFRP5* (Fig. 4e,f and Extended Data Fig. 8a). We observed several sub-populations within the organoids. *MUC6* expression varied with only some cells highly expressing *MUC6* (Fig. 4f). Furthermore, *ASGR1* was lowly expressed on the transcript level in few organoid cells (Fig. 4g), although it was clearly detected on the protein level (Fig. 3h). *SERPINA3*, *SERPINA1* and *CLU*, which are highly expressed by hepatocytes and hepatocyte progenitors, are upregulated in a non-dividing *MUC6*^-^ population from the organoids (cluster 2) (Fig. 4g, Extended Data Fig. 8b). Moreover, gene set enrichment analysis (GSEA) shows that these cells upregulate hepatocyte pathways, e.g. IGF transport and uptake by IGFBPs, and downregulate WNT signaling and cell cycle genes (Fig. 4h). *CFTR* expression marks another group of non-dividing *MUC6*^-^ cells in the organoids, corresponding to the cholangiocyte lineage (Fig. 4g). Hence, *SERPINA3*^+^ and *CFTR*^+^ organoid populations likely correspond to more mature differentiating hepatocyte and cholangiocyte progenitors, respectively, while the remaining population comprises many *MKI67*^+^ dividing cells at different stages of the cell cycle (Extended Data Fig. 8c).

Our comparison between the patients’ cells and organoid cells reveals many differentially expressed genes and pathways, e.g. upregulation of WNT signaling and cell cycle in organoids (Fig. 4e,i). Organoid cells express *AGR2* and other mucin family genes like *MUC5AC* and *MUC5B*, which are not expressed by the patients’ cells but are expressed in intestinal cells and gastrointestinal cancers^42-45^ (Extended Data Fig. 8b). Furthermore, cells from the organoids drastically downregulate *ALB* and express *AKR1B10*, a marker of hepatocellular carcinoma^46,47^ (Extended Data Fig. 8a,c). These observations reflect that EPCAM^+^ cells within the organoids acquire a more proliferative state and appear to upregulate stem cell related pathways like WNT signaling which are also active in *EPCAM*^+^*MUC6*^+^ liver cells.

Together with these functional validation experiments, the gene signature of the *MUC6*^+^ cells, comprising low levels of cholangiocyte genes, the in situ location, as well as the slightly increased cholangiocyte bias inferred by FateID suggest that these cells can be defined as a population of bile duct cells with a putative stem cell nature.

### The human liver cell atlas as a reference for identifying perturbed cellular phenotypes in hepatocellular carcinoma

Hepatocellular carcinoma is the most common type of primary liver cancer, usually occurring in people with chronic liver diseases like cirrhosis^48^. Prevalence, incidence, morbidity and mortality of HCC have been rising due to unsatisfactory treatment options and increasing underlying causes such as obesity and metabolic disorders^3^. In order to demonstrate the value of our atlas as a reference for comparing with cells from diseased livers, we sorted and sequenced CD45^+^ and CD45^-^ single cells from HCC tissue surgically resected from three different patients (Extended Data Fig. 9a,b, Methods).

We were able to recover several cell types from the tumors, including cancer cells, endothelial cells, Kupffer cells, NKT and NK cells (Fig. 5a,b). We then compared each of the corresponding cell types from the HCC tumors with those from the normal liver cell atlas. Differential gene expression analysis and GSEA revealed that cancer cells from HCC down-regulate hepatocyte genes like *ALB* and *PCK1*, and lose the metabolic signature of normal hepatocytes (Fig. 5c,d and g). In contrast, they express *AKR1B10*, a known biomarker of HCC with potential involvement in hepatocellular carcinogenesis^46,47^ (Fig. 5c). Moreover, some cancer cells expressed *CD24*, a cancer stem cell marker^49^ that was also expressed in normal *EPCAM*^+^ liver cells (Fig. 5c). We found that cancer cells upregulate WNT and Hedgehog signaling pathways, which are active in *EPCAM*^+^ cells relative to normal hepatocytes, highlighting similarities between *EPCAM*^+^ normal liver progenitors and the observed cancer cell population (Fig. 5d,g). Unexpectedly, expression of known pan-HCC markers like *S100A9* and *NTS* was restricted to a subpopulation of cancer cells. Cancer cells with low expression of these markers upregulated *LTB*, *PKM*, and *TRAC* (Fig. 5d). Similarly, the pan-HCC marker *GPC3* was not expressed in all cancer cells but upregulated in stellate cells from the tumor (Fig. 5c).

**Figure 5.**
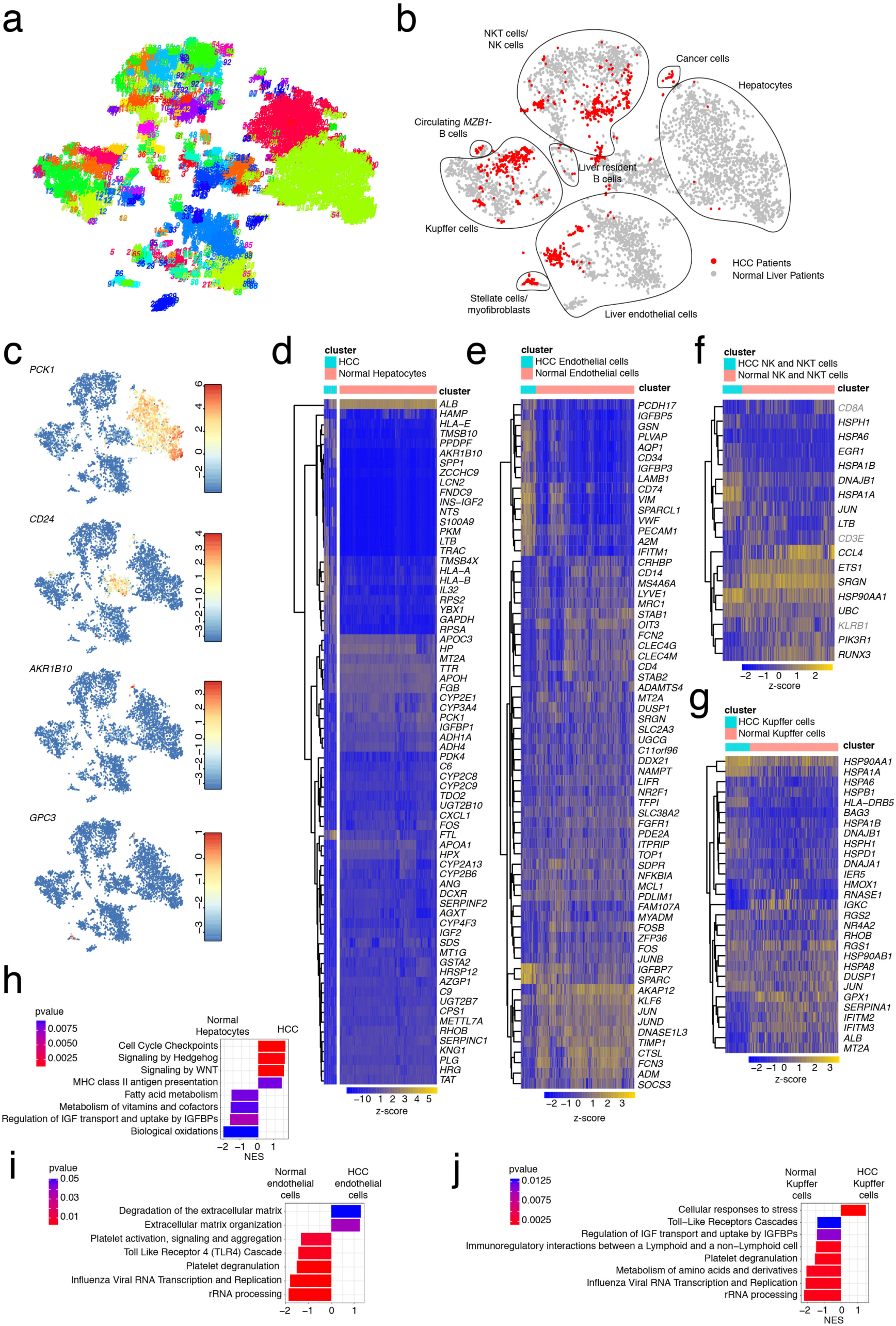
ScRNA-seq of patient-derived HCC reveals cancer-specific genes signatures and perturbed cellular phenotypes. **a**, t-SNE map showing RaceID3 clusters for normal liver cells co-analyzed with cells from HCC tissues of three different patients. **b**, Symbol t-SNE map highlighting normal liver cells and cells from HCC. The main cell types are labeled. **c**, Expression t-SNE maps of *PCK1*, *CD24*, *AKR1B10* and *GPC3*. **d**, Heatmap of differentially expressed genes between cancer cells from HCC and normal hepatocytes. Differentially expressed genes were inferred by pairwise comparison between cancer cells from HCC clusters and normal cells from hepatocyte clusters (Benjamini-Hochberg corrected *P*<0.05) **e**, Heatmap of differentially expressed genes between endothelial cells from HCC and normal endothelial cells. Differentially expressed genes were inferred by pairwise comparison between HCC endothelial cells from HCC clusters and normal endothelial cells from MaVEC and LSEC clusters (Benjamini-Hochberg corrected *P*<0.05). **f**, Heatmap of differentially expressed genes between NK and NKT cells from HCC and normal NK and NKT cells. Differentially expressed genes were inferred by pairwise comparison between HCC NK and NKT cells from HCC clusters and normal cells from NK and NKT cell clusters (Benjamini-Hochberg corrected *P*<0.05). **g**, Heatmap of differentially expressed genes between Kupffer cells from HCC and normal Kupffer cells. Differentially expressed genes were inferred by pairwise comparison between HCC Kupffer cells from HCC clusters and normal cells from Kupffer cell clusters (Benjamini-Hochberg corrected *P*<0.05). **h**, GSEA for differentially expressed genes between normal hepatocytes and cancer cells from HCC. **i**, GSEA for differentially expressed genes between normal endothelial cells and endothelial cells from HCC. **j**, GSEA for differentially expressed genes between normal Kupffer cells and Kupffer cells from HCC. The bar charts in (h), (i), and (j) show the normalized enrichment score (NES) and highlight the p-value.

Endothelial cells from the tumor up-regulate the expression of extracellular matrix organization genes such as *COL4A1*, *COL4A2* and *SPARC* and insulin growth factor binding protein genes like *IGFBP7* in comparison to normal endothelial cells (Fig. 5e,h and Extended Data Fig. 9c,d).

Immune cell populations from the tumor, comprising Kupffer cells, NKT and NK cells upregulate stress response genes and lose their immunoregulatory functions compared to their normal liver counterparts (Fig. 5f,g,h and Extended Data Fig. 9e). Moreover, Kupffer cells and endothelial cells from the tumor lose their subsets and zonation signatures found in normal liver (Fig. 5h,I and Extended Data Fig. 9c). We conclude that the comparison of scRNA-seq data between the cell populations of HCC and the liver cell atlas allows the inference of perturbed gene expression signatures and modulated functions across cell types.

### Investigating perturbed cellular phenotypes in a human liver chimeric mouse model

Patient derived xenograft mouse models, or humanized mouse models, are a powerful tool for studying human cells and diseases *in vivo*. Human liver chimeric mice have been developed as state-of-the-art models^50,51^. In these models, human hepatocytes are transplanted into the mouse liver by intra-splenic injection. While the engrafted human hepatocytes are similar to their counterparts in the human liver as shown by histology, biological profile and susceptibility to infection with human tropic pathogens, the transcriptional state of the engrafted cells is largely unknown. To correctly interpret such experiments, it is crucial to understand and take into account the differences between cells directly taken from the human liver and transplanted human cells in the mouse model. It can be expected that both the transplantation procedure itself and the mouse microenvironment may affect the state of these cells, leading to conclusions which are potentially valid only in the humanized mouse model and not in the human.

To address this issue, we transplanted human liver cells from patient-derived hepatocyte and non-parenchymal cell fractions in an *FAH*^-/-^ mouse liver model^50,52^ and sorted single human cells in an unbiased fashion and on the basis of hepatocyte and endothelial cell markers post engraftment for scRNA-seq (Fig. 6a). We then used our liver cell atlas to compare human cells engrafted into the mouse model to cells sequenced directly from the patients and observed that we had successfully transplanted both human hepatocytes and endothelial cells in the *FAH*^-/-^ mouse liver model (Fig. 6b-f). Surprisingly, engrafted human (HMouse) hepatocytes and endothelial cells mostly cluster separately from their human liver counterparts (Fig. 6b). Both transplanted cell types maintained their fundamental gene signatures, such as *ALB* and *PCK1* expression by HMouse hepatocytes (Fig. 6d) and *CLEC4G* or *CD34* expression by HMouse endothelial cells (Fig. 6f), but nevertheless express many genes differentially from human liver cells (Fig. 6e,g and Extended Data Fig. 10). Specifically, we observed genes that were only expressed by the engrafted cells such as *AKR1B10*, which was also expressed by cancer cells from HCC and *EPCAM*^+^ cells from the organoids but not in the normal human liver cells (Fig. 6e). Vice versa, a number of genes, e.g. *CXCL1*, were expressed by hepatocytes from the human liver but not by HMouse hepatocytes (Fig. 6e). Other genes were expressed in both human liver and transplanted cells yet at substantially different levels, e.g. *FTL* or *HP* (Fig. 6g, Extended Data Fig. 10). GSEA of differentially expressed genes revealed that HMouse hepatocytes and endothelial cells downregulated zonated pathways, such as regulation of IGF transport and uptake by IGFBPs, and upregulated WNT and Hedgehog signaling as well as cell cycle genes (Fig. 6h). Notably, these were the same pathways that we found to be upregulated in the HCC cancer cells, EPCAM^+^ liver progenitors from the patients and the organoids.

**Figure 6.**
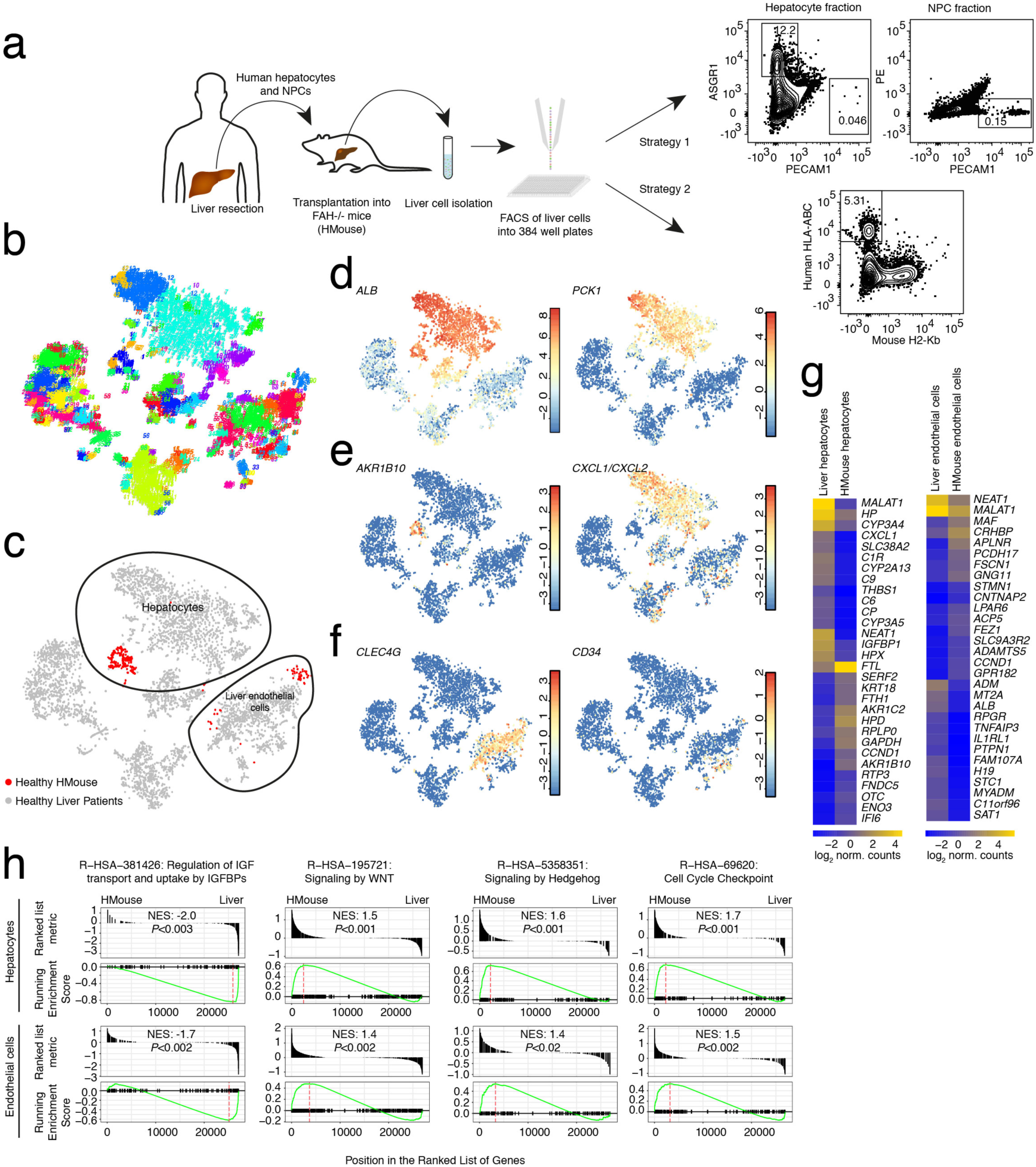
Exploring the gene expression signature of human liver cells in a humanized mouse model. **a**, Outline of the transplantation of human liver cells (hepatocytes and non-parenchymal cells) into the *FAH*^-/-^ mouse and the two strategies used to sort human cells from the mouse liver for scRNA-seq. **b**, t-SNE map of RaceID3 clusters of liver cells from the patients co-analyzed with cells from the humanized mouse liver model. **c**, Symbol t-SNE map highlighting normal liver cells and cells from the humanized mouse model. The main engrafted cell types (hepatocytes and endothelial cells) are circled. **d**, Expression t-SNE maps of hepatocyte markers *ALB* and *PCK1*. **e**, Expression t-SNE maps of *AKR1B10* and *CXCL1*/*CXCL2.* **f**, Expression t-SNE maps of endothelial markers *CLEC4G* and *CD34*. **g**, Heatmaps of differentially expressed genes between hepatocytes and endothelial cells from the patients (Liver hepatocytes and Liver endothelial cells) and from the humanized mouse model (HMouse hepatocytes and HMouse endothelial cells). Differentially expressed genes were inferred by pairwise comparisons between the HMouse clusters and the patient clusters for each cell type (Benjamini-Hochberg corrected *P*<0.01). The 15 most upregulated and 15 most downregulated genes are included. **h**, GSEA of differentially expressed genes between hepatocytes and endothelial cells from the humanized mouse (HMouse) and from the patients (Liver). Differentially expressed genes were inferred as described in (g).

Such progenitor gene signatures are likely established to enable successful engraftment requiring proliferation of transplanted cells in the mouse liver. Other gene signatures, such as those related to metabolic zonation, likely change in response to the mouse liver microenvironment, which is different from the human counterpart. In summary, these data reveal that mouse transplanted human liver cells retain their fundamental gene signature but revert to an immature phenotype upon transplantation.

## DISCUSSION

We here established a human liver cell atlas, comprising all major cell types in the adult liver and revealing heterogeneity within these populations. Being able to perform scRNA-seq on cryopreserved liver cells broadens the application of this approach to material from biobanks and avoids logistical difficulties associated with the retrieval of fresh human tissue from liver biopsies or surgical resections.

A potential weakness is that liver tissue was obtained from hospitalized patients. Hence, interference of medication, surgical procedure (such as ischemia) or the presence of other diseases outside the resected normal tissue (e. g. colorectal cancer, cholangiocarcinoma) on liver cell gene expression cannot be excluded. However, for ethical and regulatory reasons it is virtually impossible to obtain liver tissue from normal volunteers without any co-morbidity since it requires an invasive intervention step.

Our atlas reveals that hepatocytes, Kupffer cells and LSECs share co-zonated pathways and genetic networks, suggesting that they cooperate in order to carry out essential functions. Further work should aim at understanding the molecular regulation, coordination and role of these liver cell subtypes and their zonation in the context of health and disease.

The existence of distinct cell types in the EPCAM^+^ compartment of the human liver raises the question of which cell types proliferate and differentiate to distinct cell lineages and contribute to liver regeneration after different types of liver damage. The novel EPCAM^+^MUC6^+^ putative naïve stem cell population provides a strong candidate with potential involvement in homeostatic turnover, liver regeneration, disease pathogenesis, and tumor formation. We anticipate that this population could be of potential use in liver regenerative medicine.

The atlas will provide a key reference to investigate liver malignancies, including metabolic, inflammatory and viral diseases as well as cancer. Our HCC proof-of-principle analysis demonstrates that the atlas enables the identification of disease-specific cellular phenotypes. We expect that such analyses could allow the discovery of novel targets for urgently needed therapeutics, such as for NASH, HCC or cholangiocarcinoma.

Finally, the methods developed in this study will contribute to advance urgently needed liver disease biology models, including organoids as well as humanized liver chimeric mouse models. Indeed, by analyzing organoids and the phenotype of engrafted human liver cells in humanized mice we unravel the similarities and differences of these models compared to the human liver.

In light of the above, our human liver cell atlas provides a powerful and innovative resource enabling the discovery of previously unknown cell types in the normal and diseased liver and will contribute to a better understanding of *in vitro* and *in vivo* human liver models.

## Acknowledgments

This study was supported by the Max Planck Society, the German Research Foundation (DFG) (SPP1937 GR4980/1-1, GR4980/3-1, and GRK2344 MeInBio), by the DFG under Germany’s Excellence Strategy (CIBSS – EXC-2189 – Project ID 390939984), and by the Behrens-Weise-Foundation (all to D.G.). This work was supported by ARC, Paris and Institut Hospitalo-Universitaire, Strasbourg (TheraHCC and TheraHCC2.0 IHUARC IHU201301187 and IHUARC2019 to T.F.B.), the European Union (ERC-AdG-2014-671231-HEPCIR to T.F.B., EU H2020-667273-HEPCAR to T.F.B, ERC-PoC-2016-PRELICAN to T.F.B), ANRS and the Foundation of the University of Strasbourg. This work was done under the framework of the LABEX ANR-10-LABX-0028_HEPSYS and Inserm Plan Cancer and benefits from funding from the state managed by the French National Research Agency as part of the Investments for the future. We thank Sebastian Hobitz and Konrad Schuldes from the FACS facility and Dr. Ulrike Bönisch from the Deep Sequencing facility of the Max Planck Institute of Immunobiology and Epigenetics. We acknowledge the CRB (Centre de Ressources Biologiques-Biological Resource Centre of the Strasbourg University Hospitals) for the management of regulatory requirements of patient-derived liver tissue. We thank Dr. Catherine Fauvelle and Laura Heydmann (Inserm U1110, University of Strasbourg) for their contributions to the initial single cell isolations used in the study. We thank Drs. Frank Juehling, François Duong and Catherine Schuster (Inserm U1110, University of Strasbourg) for helpful discussions. We thank the patients for providing informed consent to participate in the study and the nurses, technicians and medical doctors of the hepatobiliary surgery and pathology services of the Strasbourg University Hospitals for their support. This publication is part of the Human Cell Atlas - http://www.humancellatlas.org/publications.

## Author contributions

T.F.B. and D.G. conceived the study. N.A. designed, optimized, and performed cell sorting, scRNA-seq experiments, organoid culture, immunofluorescence, provided validation using the Human Protein Atlas, and performed computational analysis and interpretation of the data. A.S. managed the supply of patient material, isolated single cells from patient tissue, performed animal experiments and immunofluorescence. S. contributed to scRNA-seq analyses and performed single-cell RNA-seq experiments. L.M. performed animal experiments. S.D. isolated single cells from patient tissues. P.P. performed liver resections and provided patient liver tissues. T.F.B. established the liver tissue supply pipeline and supervised the animal experiments. D.G. analyzed and interpreted the data and supervised experiments and analysis of N.A. and S. D.G., N.A., and T.F.B. coordinated and led the study. N.A. and D.G. wrote the manuscript with input from S., A.S., and T.F.B.

## Competing interests

The authors declare no competing interests.

## METHODS

### Human liver samples

Human liver tissue samples were obtained from patients who had undergone liver resections between 2015 and 2018 at the Center for Digestive and Liver Disease (Pôle Hépato-digestif) of the Strasbourg University Hospitals University of Strasbourg, France. For the human liver cell atlas, samples were acquired from patients without chronic liver disease (defined as liver damage lasting over a period of at least six months), genetic hemochromatosis with homozygote C282Y mutation, active alcohol consumption (> 20 g/d in women and > 30 g/d in men), active infectious disease, pregnancy or any contraindication for liver resection. All patients provided a written informed consent. The protocols followed the ethical principles of the declaration of Helsinki and were approved by the local Ethics Committee of the University of Strasbourg Hospitals and by the French Ministry of Education and Research (Ministère de l'Education Nationale, de l’Enseignement Supérieur et de la Recherche; approval number DC-2016-2616). Data protection was performed according to EU legislation regarding privacy and confidentiality during personal data collection and processing (Directive 95/46/EC of the European Parliament and of the Council of the 24 October 1995).

### Tissue dissociation and preparation of single cell suspensions

Human liver specimens obtained from resections were perfused for 15 minutes with calcium-free 4-(2-hydroxyethyl)-1-piperazine ethanesulfonic acid buffer containing 0.5 mM ethylene glycol tetraacetic acid (Fluka) followed by perfusion with 4-(2-hydroxyethyl)-1-piperazine ethanesulfonic acid containing 0.5 mg/mL collagenase (Sigma-Aldrich) and 0.075% CaCl_2_ at 37°C for 15 min as previously described^53^. Then the cells were washed with phosphate-buffered saline (PBS) and nonviable cells were removed by Percoll (Sigma-Aldrich) gradient centrifugation. Part of the isolated cells was further separated into primary human hepatocytes (PHH) and non-parenchymal cells (NPCs) by an additional centrifugation step. The isolated cells were frozen in liquid nitrogen using the CryoStor^®^ CS10 solution (Sigma-Aldrich).

### Transplantation of human cells into *Fah^−/−^/Rag2^−/−^/Il2rg ^−/−^* mice

*Fah^−/−^/Rag2^−/−^/Il2rg ^−/−^ (FRG*) breeding mice were kept at the Inserm Unit 1110 SPF animal facility and maintained with 16 mg/L of 2-(2-nitro-4-trifluoro-methyl-benzoyl)-1,3 cyclohexanedione (NTBC; Swedish Orphan Biovitrum) in drinking water. Six-week old mice were intravenously injected with 1.5 x 10^9^ pfu of an adenoviral vector encoding the secreted form of the human urokinase-like plasminogen activator (Ad-uPA)^54^. Forty-eight hours later, 10^6^ PHH and 2 x 10^5^ NPCs from the same liver donor and isolated as previously described, were injected intra-splenically via a 27-gauge needle. For the procedure, the mice were kept under gaseous isoflurane anesthesia and received a subcutaneous injection of buprenorphine at the dose of 0.1 mg/kg. After the transplantation the NTBC was gradually decreased and completely withdrawn in 7 days. The success of the transplantation was evaluated 2 months after the procedure by dosing human albumin in mouse serum as previously described^50^. All procedures are consistent with the guidelines set by the Panel on Euthanasia (AVMA) and the NIH Guide for the Care and Use of Laboratory Animals as well as the Declaration of Helsinki in its latest version, and to the Convention of the Council of Europe on Human Rights and Biomedicine. The animal research was performed within the regulations and conventions protecting the animals used for research purposes (Directive 86/609/EEC), as well as with European and national laws regarding work with genetically modified organs. The animal facility at the University of Strasbourg, Inserm U1110 has been approved by the regional government (Préfecture) and granted the authorization number N° D67-482-7, 2012/08/22.

### FACS

Liver cells were sorted from mixed, hepatocyte, and non-parenchymal cell fractions on an Aria Fusion I using a 100 μm nozzle. Cells from the HCC samples were not fractionated and were sorted directly after tissue digestion. Zombie Green (Biolegend) was used as a viability dye. Cells were stained with human specific antibodies against CD45 (Biolegend), PECAM1 (Biolgend), CD34 (Biolegend), CLEC4G (R&D systems), ASGR1 (BD Biosciences), EPCAM (R&D systems), CFTR (LSBio) and TROP2 (Biolegend). Organoids were stained with antibodies against EPCAM and ASGR1. For the humanized mouse samples, cells were stained either with antibodies against ASGR1 and PECAM1 or human HLA-ABC (BD Biosciences) and mouse H2-Kb (BD Biosciences). Viable cells were sorted in an unbiased fashion or from specific populations based on the expression of markers into the wells of 384 well plates containing lysis buffer.

### Single-cell RNA amplification and library preparation

Single cell RNA sequencing was performed according to the mCEL-Seq2 protocol^7^. Viable liver cells were sorted into 384-well plates containing 240 nL of primer mix and 1.2 μL of PCR encapsulation barrier, Vapor-Lock (QIAGEN) or mineral oil (Sigma-Aldrich). Sorted plates were centrifuged at 2200 g for a few minutes at 4°C, snap-frozen in liquid nitrogen and stored at −80°C until processed. 160nL of reverse transcription reaction mix and 2.2 μL of second strand reaction mix were used to convert RNA into cDNA. cDNA from 96 cells were pooled together before clean up and in vitro transcription, generating 4 libraries from one 384-well plate. 0.8 μL of AMPure/RNAClean XP beads (Beckman Coulter) per 1 μL of sample were used during all the purification steps including the library cleanup. Other steps were performed as described in the protocol. Libraries were sequenced on an Illumina HiSeq 2500 and 3000 sequencing system (pair-end multiplexing run, high output mode) at a depth of ~150,000-200,000 reads per cell.

### Quantification of Transcript Abundance

Paired end reads were aligned to the transcriptome using bwa (version 0.6.2-r126) with default parameters^55^. The transcriptome contained all gene models based on the human whole genome ENCODE V24 release. All isoforms of the same gene were merged to a single gene locus. The right mate of each read pair was mapped to the ensemble of all gene loci and to the set of 92 ERCC spike-ins in the sense direction. Reads mapping to multiple loci were discarded. The left read contains the barcode information: the first six bases corresponded to the cell specific barcode followed by six bases representing the unique molecular identifier (UMI). The remainder of the left read contains a polyT stretch. The left read was not used for quantification. For each cell barcode, the number of UMIs per transcript was counted and aggregated across all transcripts derived from the same gene locus. Based on binomial statistics, the number of observed UMIs was converted into transcript counts^56^.

### Single-Cell RNA Sequencing Data Analysis

5416 cells passed quality control thresholds for the normal human liver cell atlas. For cells from the organoids, 625 cells passed the quality control thresholds. For cells from HCC, 911 cells passed the quality control thresholds. For cells from the humanized mouse, 315 cells passed the quality control threshold. All the datasets were analyzed using RaceID3^10^. Rescaling to 1,200 transcripts per cell was applied for data normalization. Prior to normalization, cells expressing >2% of *Kcnq1ot1* transcripts, a previously identified marker of low quality cells were removed from the analysis. Moreover, transcripts correlating to *Kcnq1ot1* with a Pearson’s correlation coefficient >0.65 were also removed. RaceID3 was run with the following parameters: mintotal=1200, minexpr=5, outminc=5, FSelect=TRUE, probthr=10^-3^. The t-distributed stochastic neighbor embedding (t-SNE) algorithm was used for dimensional reduction and cell cluster visualization.

### Diffusion Pseudo-Time Analysis and Self-Organizing Maps

Diffusion pseudotime (dpt) analysis^28^ was implemented and diffusion maps generated using the destiny R package. The number of nearest neighbors, k, was set to 100. SOMs were generated using the FateIDpackage based the ordering computed by dpt as input.

### Lineage Analysis of the *EPCAM*^+^ compartment

For a separate analysis of the *EPCAM*^+^ population, all cells from clusters 13, 17, 45, 48, and 59 were extracted and reanalyzed with RaceID3^10^ using the parameters mintotal=1200, minexpr=2, outminc=2, probthr=1e-4 and default parameters otherwise. StemID2^10^ was run on these clusters with cthr=5 and nmode=TRUE. FateID^10^ was run on the filtered and feature-selected expression matrix from RaceID3, with target clusters inferred by FateID using *ASGR1* and *CXCL8* as markers for hepatocyte and cholangiocyte lineage target clusters.

### Differential Gene Expression Analysis

Differential gene expression analysis between cells and clusters was performed using the diffexpnb function from the FateID package.

### Pathway Enrichment Analysis and Gene Set Enrichment Analysis

Symbol gene IDs were first converted to Entrez gene IDs using the clusterProfiler^57^ package. Pathway enrichment analysis and GSEA^58,59^ were implemented using the ReactomePA^60^ package. Pathway enrichment analysis was done on genes taken from the different modules in the SOMs. GSEA was done using the differentially expressed genes inferred by the diffexpnb function from the FateID package without filtering on lowly expressed genes.

### Immunofluorescence

Human liver tissue was fixed in 3.7% formaldehyde. The tissue was then embedded in OCT and kept at −80°C. The tissue was cryosectioned into 5 micron sections. The tissue was washed twice for 5 min in 0.025% Triton 1x TBS (Tris-Buffered-Saline). The tissue was then blocked in 10% FBS with 1% BSA in 1x TBS for 2 hours at room temperature. The dilution used for the anti-human MUC6 FITC conjugated antibody (MBS2055834) was 1:50 in 100 μL of 1xTBS with 1% BSA. The antibody was incubated overnight at 4°C in the dark. The tissue was then washed twice with 0.025% Triton 1x TBS, dried with a tissue. After that, DAPI Fluoromount-G (Southern Biotech) was added to the tissue and coverslip placed on top. Imaging was done using the Zeiss confocal microscope LSM780. Images were taken at 20x magnification.

### Organoid culturing

Organoid culturing was done as previously described^61^. The cell populations from the EPCAM+ compartment were sorted on an Aria Fusion 1 using a 100 μm nozzle into tubes containing culture medium supplemented with 10 □M ROCK inhibitor (Y-27632) (Sigma-Aldrich). After sorting, cells were centrifuged in order to remove the medium and then resuspended in 25 μL of Matrigel. Droplets of the Matrigel solution containing the cells were added to the wells of a 24 well suspension plate and incubated for 5-10 min at 37°C until the Matrigel solidified. Droplets were overlaid with 250 μL of liver isolation medium and then incubated at 37°C, 5% CO_2_. After 3-4 days, the liver isolation medium was replaced with liver expansion medium. Organoids were passaged 13 days after isolation and then passaged multiple times 5-7 days after splitting. For FACS, single cell suspensions were prepared from the organoids by mechanical dissociation followed by TrypLE (Life Technologies) digestion as previously described^61^.

### Data availability

Data generated during this study have been deposited in Gene Expression Omnibus (GEO) with the accession code GSEXXX.

## EXTENDED DATA FIGURES

**Extended Data Figure 1.**
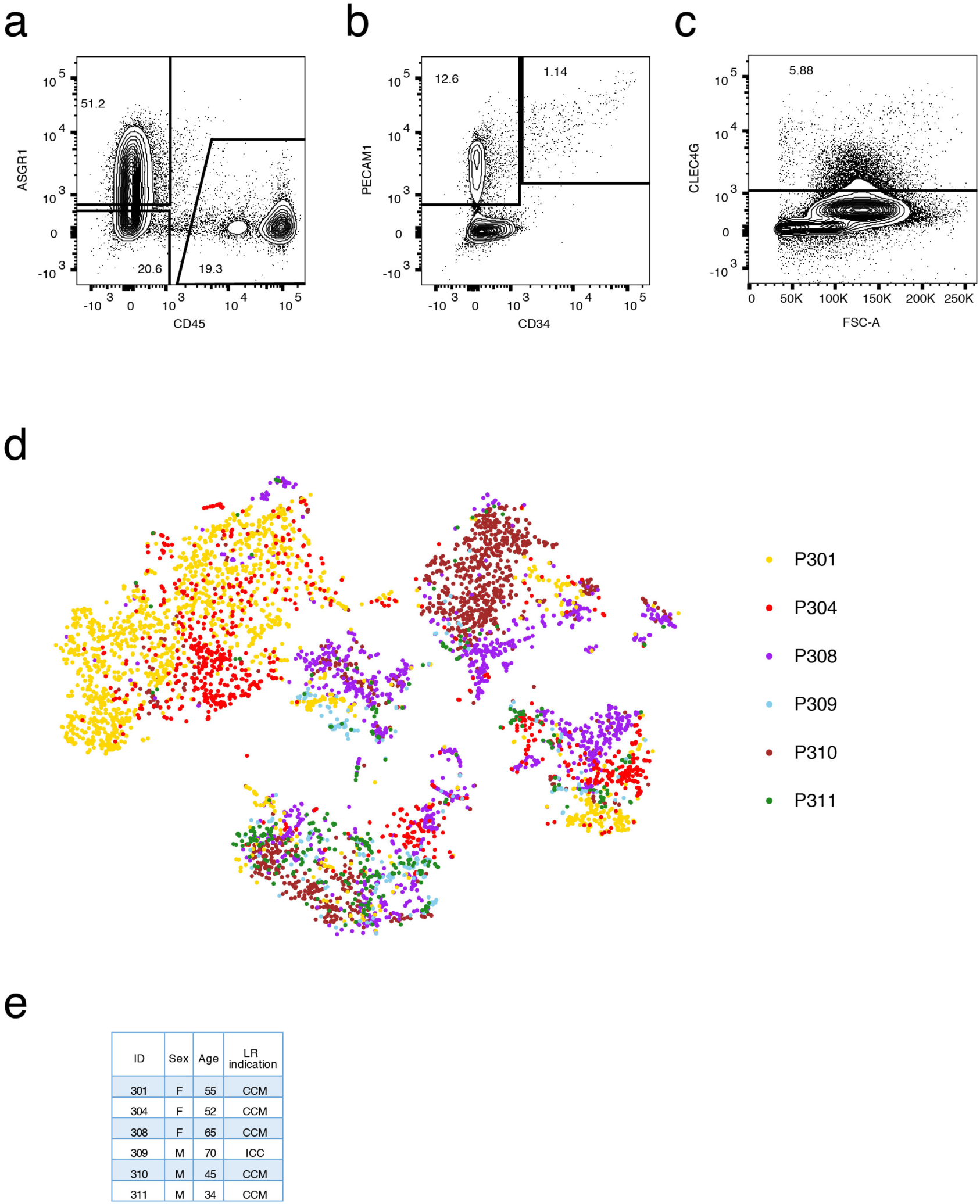
ScRNA-seq analysis of normal liver resection specimens from six adult patients. **a**, FACS plot for CD45 and ASGR1 staining from a mixed fraction (hepatocyte and non-parenchymal cells). **b**, FACS plot for PECAM1 and CD34 staining from a mixed fraction. **c**, FACS plot for CLEC4G staining from a mixed fraction. **d**, t-SNE map showing the IDs of the 6 patients from which the cells were sequenced. Cells were sequenced from freshly prepared single cell suspensions for P301 and P304 and from cryopreserved single cell suspensions for P301, P304, P308, P309, P310 and P311. Cells were sorted and sequenced mainly in an unbiased fashion from non-parenchymal cell, hepatocyte and mixed fractions of patients P301 and P304. Non-parenchymal and mixed fractions were used to sort for specific populations on the basis of markers. CD45 negative and positive cells were sorted from all patients. CLEC4G^+^ LSECs were sorted by FACS from patient P310. EPCAM positive cells were sorted by FACS from patients P308, P309, P310 and P311. **e**, Table showing patient information. CCM: colon cancer metastasis; ICC: intrahepatic cholangiocarcinoma; LR: liver resection.

**Extended Data Figure 2.**
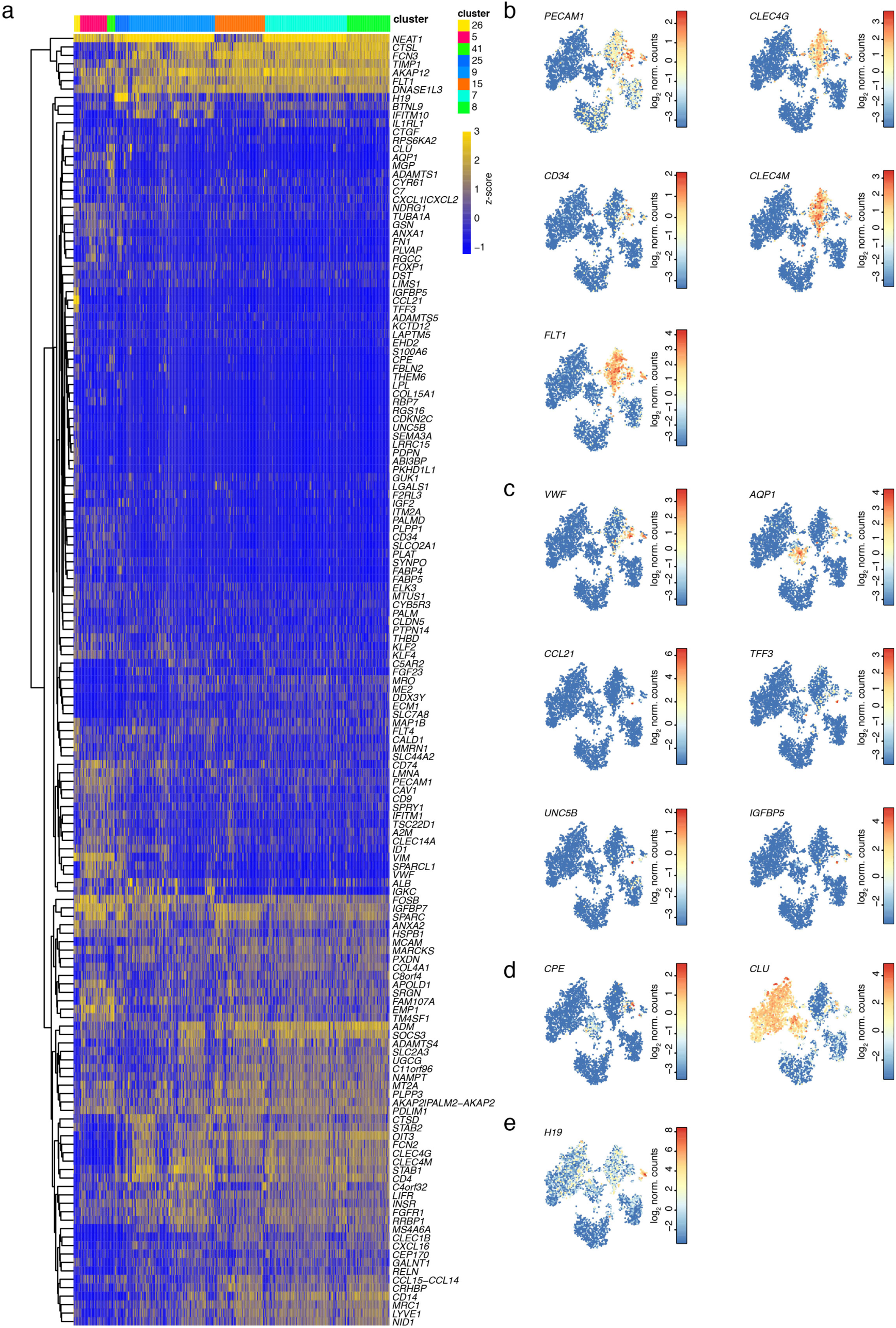
The endothelial cell compartment is a heterogeneous mixture of sub-populations. **a**, Heatmap showing differentially expressed genes between endothelial cell clusters. **b**, Expression t-SNE maps for LSEC and MaVEC markers *PECAM1*, *CLEC4G*, *CD34*, *CLEC4M* and *FLT1*. **c**, Expression t-SNE maps for *VWF*, *AQP1*, *CCL21*, *TFF3* and *UNC5B* and *IGFBP5*. **d**, Expression t-SNE maps for *CPE* and *CLU*. **e**, Expression t-SNE map for *H19*.

**Extended Data Figure 3.**
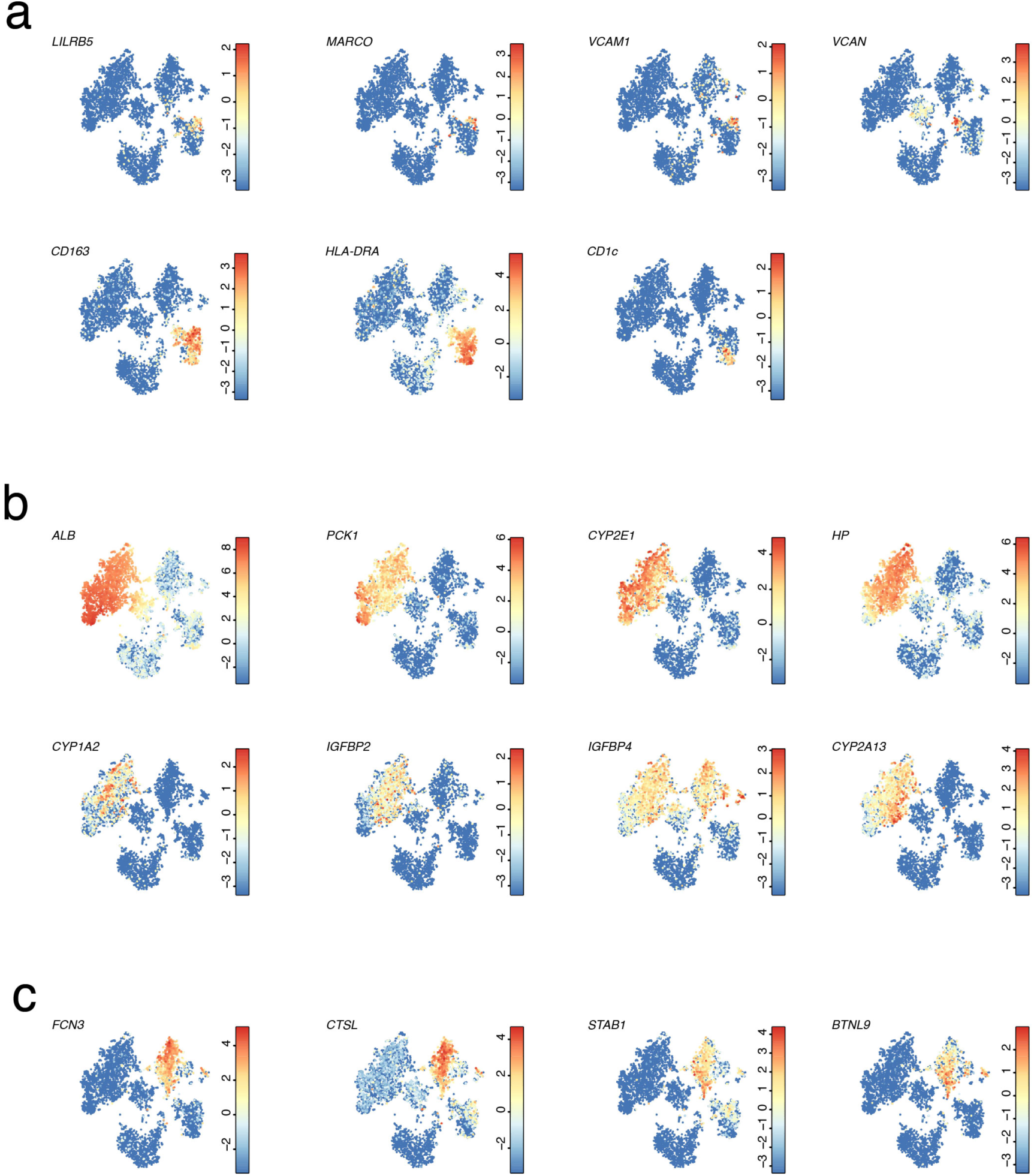
Zonated gene signatures in major human liver cell types. **a**, Expression t-SNE maps of the markers for the Kupffer cell subtypes *LILRB5*, *MARCO*, *VCAM1*, *VCAN*, *CD163*, *HLA-DRA*, and *CD1C*. **b**, Expression t-SNE maps of the zonated hepatocyte genes *ALB*, *PCK1*, *CYP2E1*, *HP*, *CYP1A2*, *IGFBP2*, *IGFBP4*, and *CYP2A13*. **c**, Expression t-SNE maps of the zonated LSEC genes *FCN3*, *CTSL*, *STAB1*, and *BTNL9*.

**Extended Data Figure 4.**
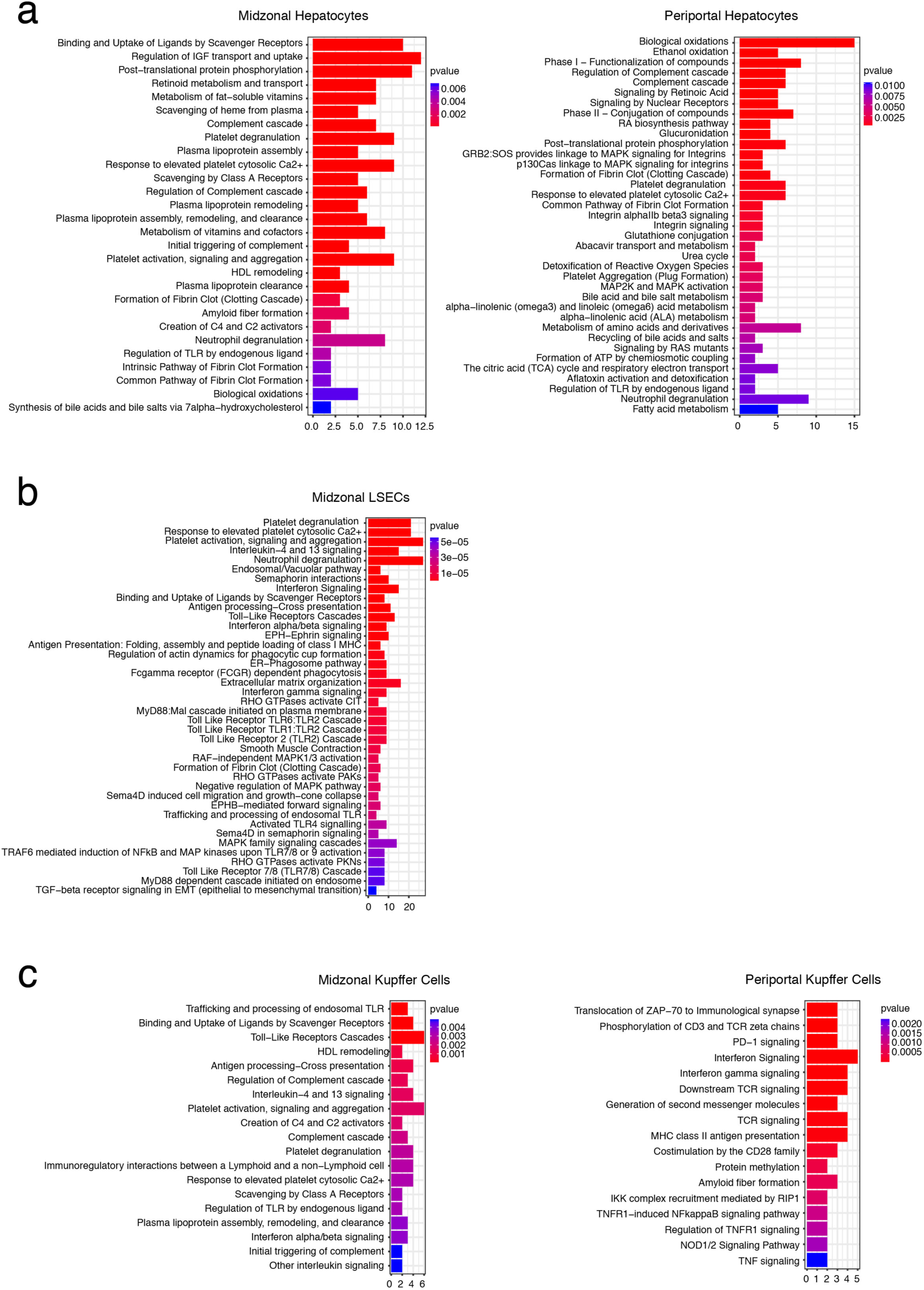
Zonated gene modules show specific enrichment of molecular pathways. **a**, Reactome pathway enrichment analysis for genes from the midzonal and periportal hepatocyte modules. **b**, Reactome pathway enrichment analysis for genes from the midzonal LSEC module. **c**, Reactome pathway enrichment analysis for genes from the midzonal Kupffer cell and periportal Kupffer cell modules. The bar charts in (a), (b), and (c) show the enrichment and highlight the p-value.

**Extended Data Figure 5.**
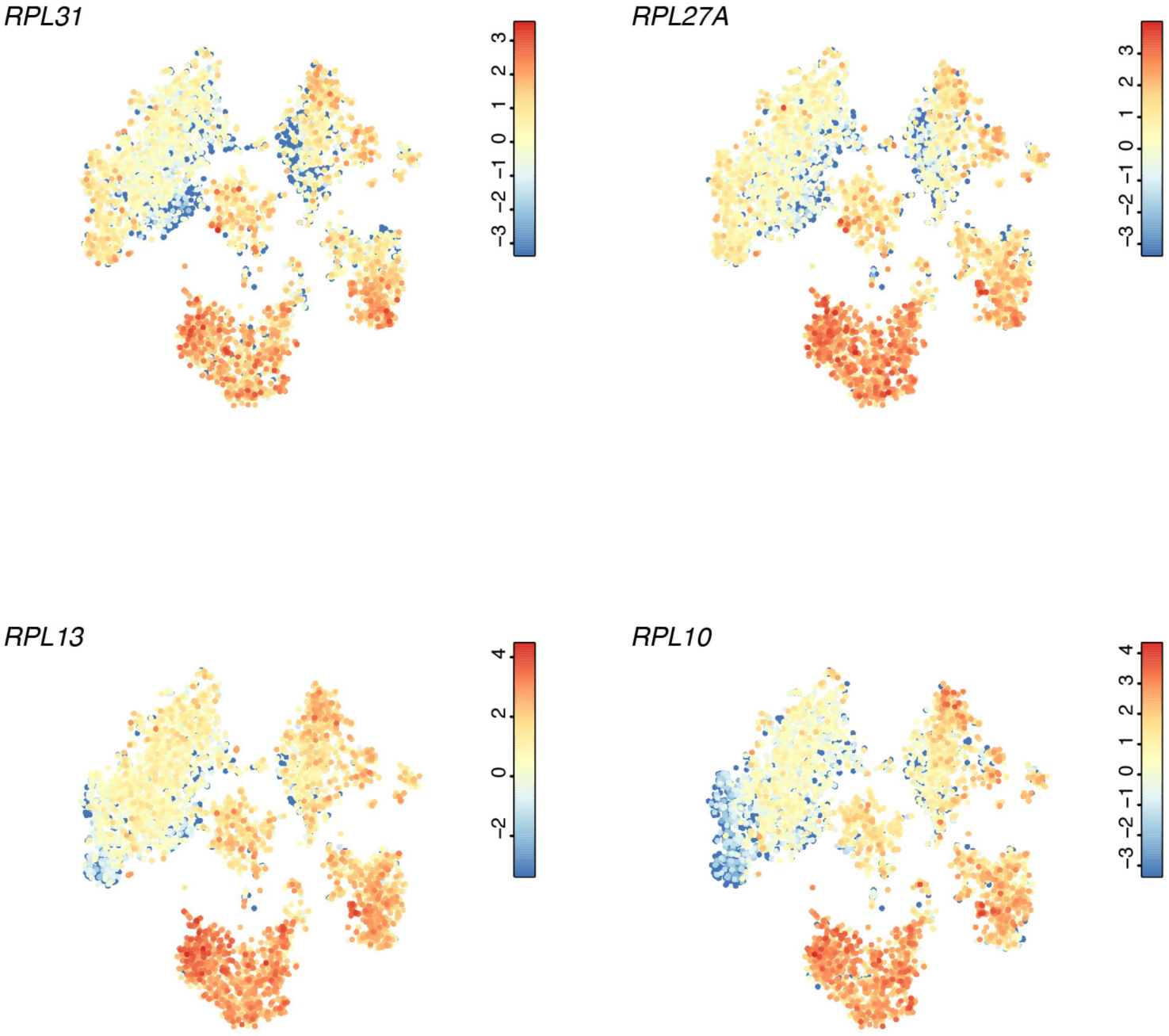
Zonated expression of ribosomal genes. Expression t-SNE maps of ribosomal genes showing their potential co-zonated expression across hepatocytes, Kupffer cells and LSECs.

**Extended Data Figure 6.**
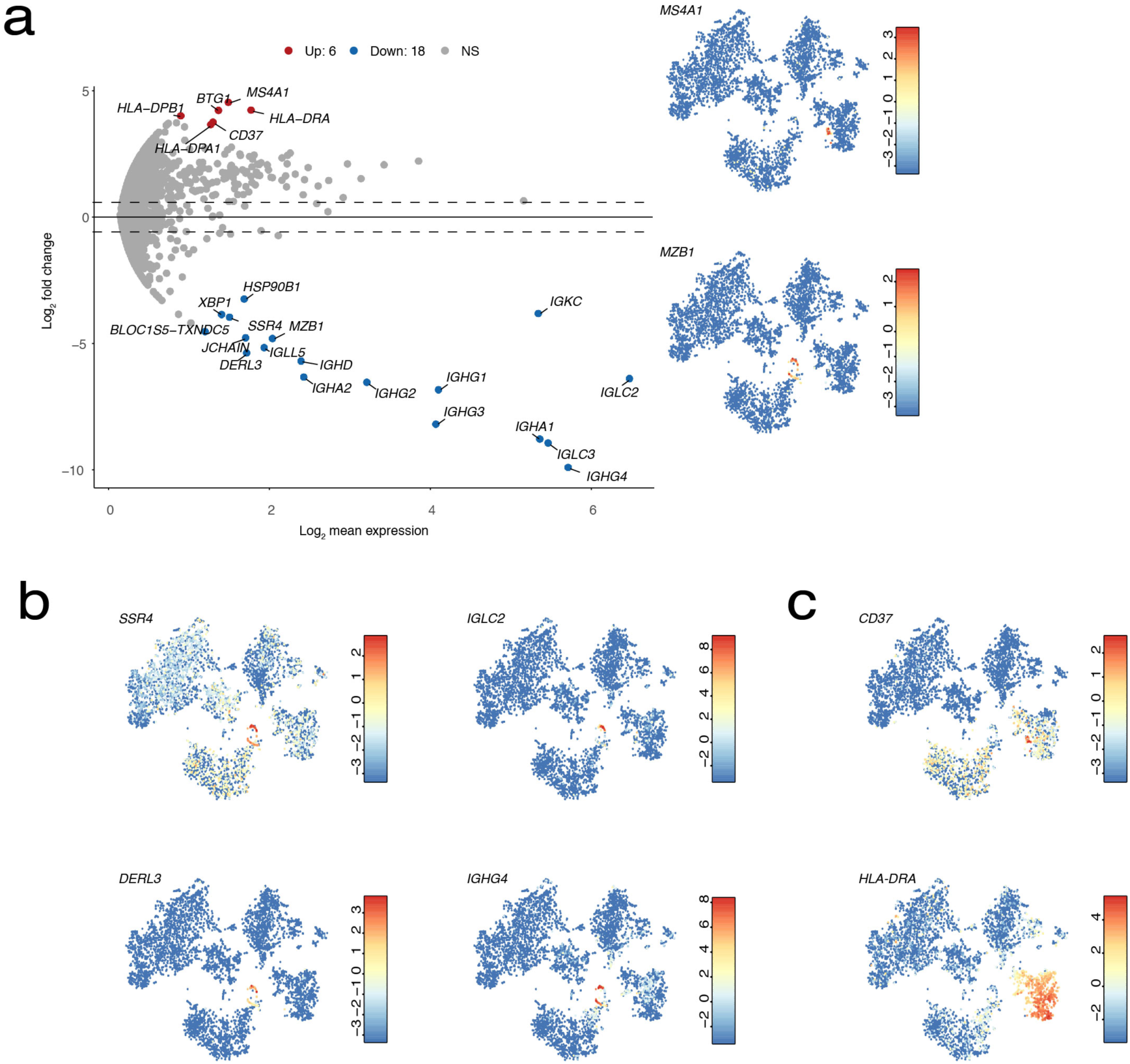
The human liver contains different B cell populations. **a**, Differentially expressed genes between *MS4A1*^+^ circulating B cells and *MZB1*^+^ liver resident B cells (Benjamini-Hochberg corrected *P*<0.05). **b**, Expression t-SNE maps of the MZB1^+^ circulating B cell-specific genes *SSR4*, *IGLC2*, *DERL3*, *IGHG4* and the *MS4A1*^+^ liver resident B cell-specific genes *CD37* and *HLA-DRA*.

**Extended Data Figure 7.**
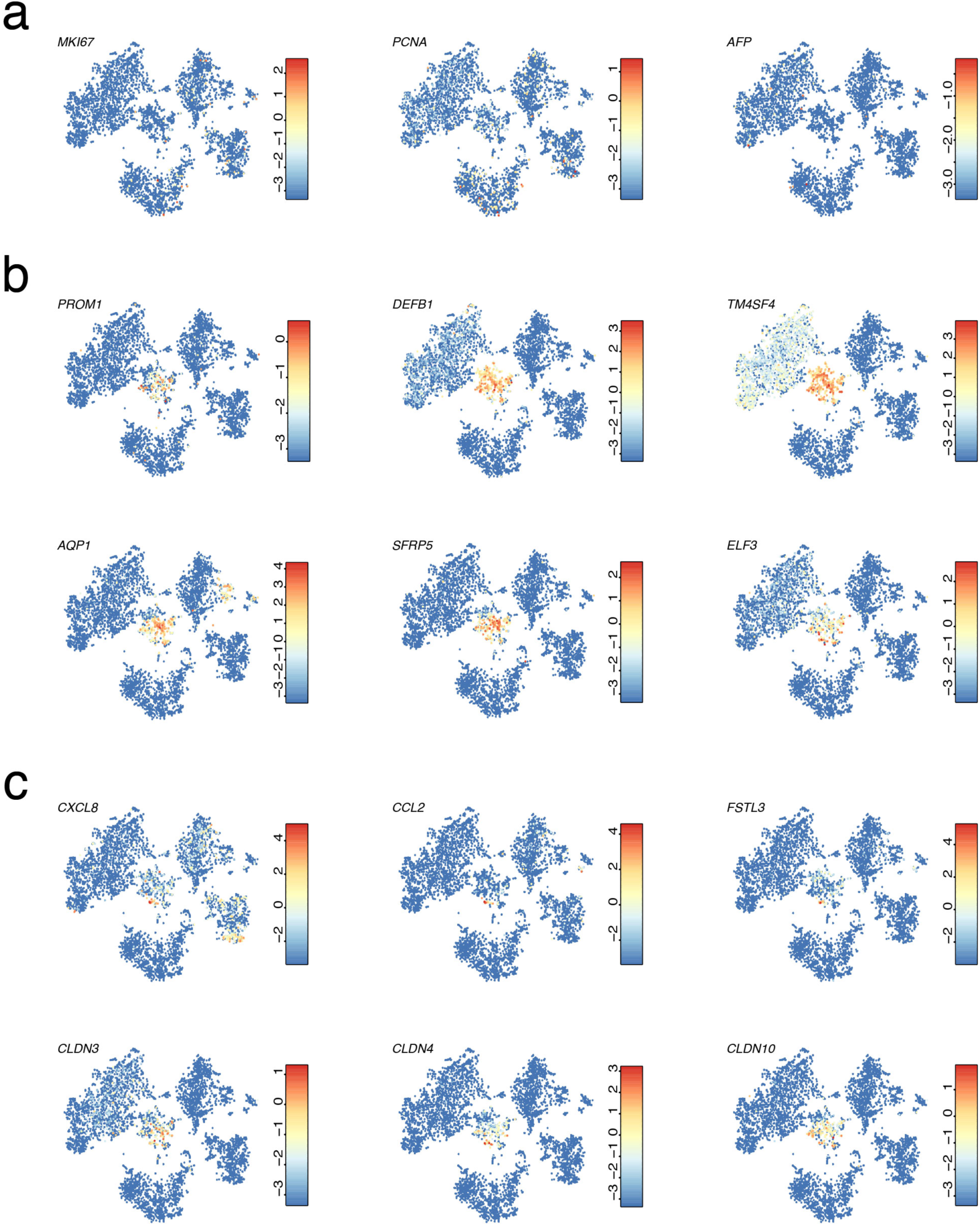
The EPCAM^+^ compartment segregates into three major sub-populations. **a**, Expression t-SNE maps of the proliferation marker *MKI67* and *PCNA* and the hepatoblast marker *AFP*. **b**, Expression t-SNE maps of EPCAM^+^ cell markers *PROM1*, *DEFB1*, *TM4SF4*, *AQP1*, *SFRP5*, and *ELF3*. **c**, Expression t-SNE maps of the cholangiocyte genes *CXCL8*, *CCL2*, *FTL3* and Claudin genes.

**Extended Data Figure 8.**
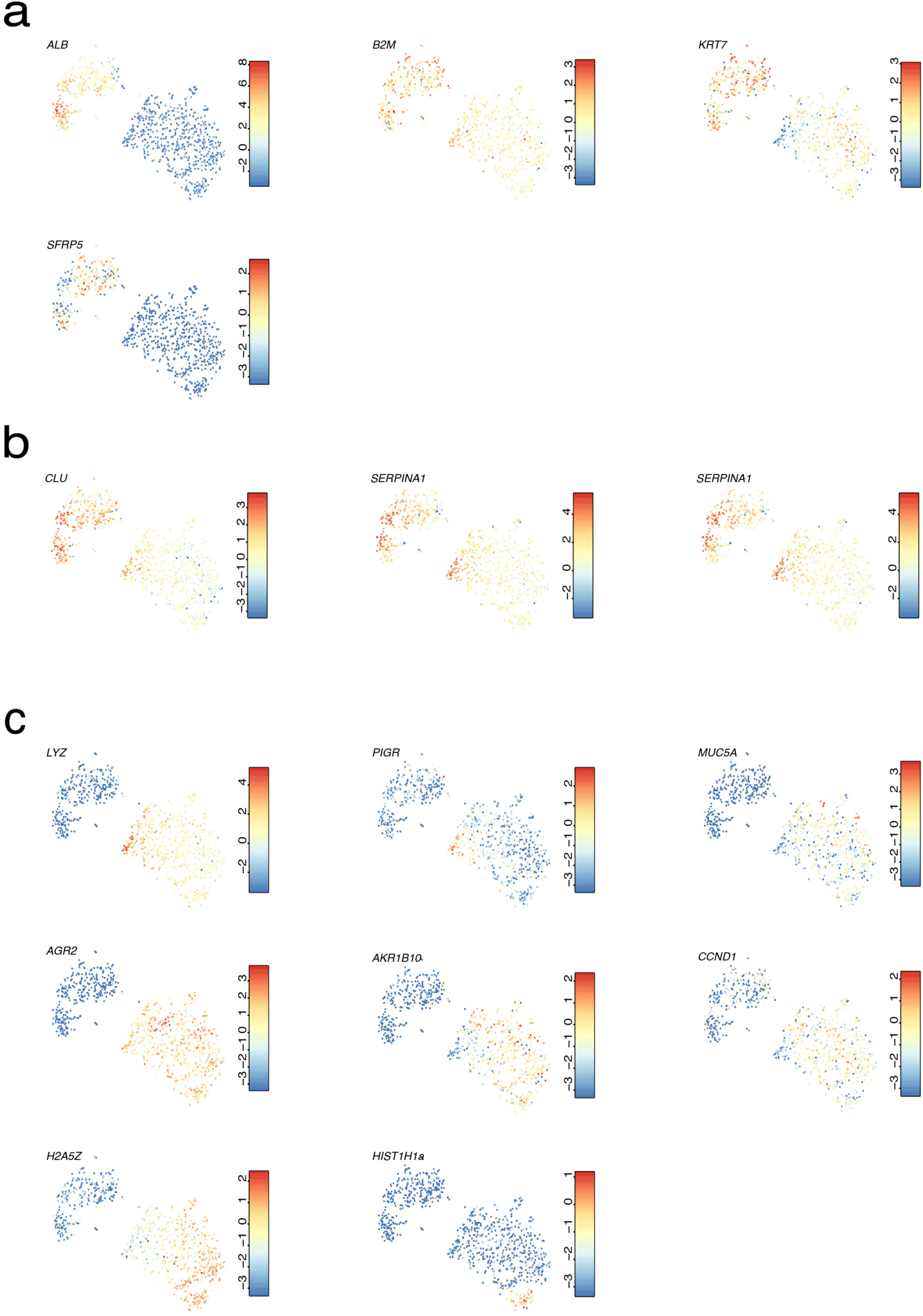
Organoid cells show a distinct gene signature in comparison to EPCAM^+^ liver cells. **a**, Expression t-SNE maps of *ALB*, *B2M*, *KRT7* and *SFRP5*. **b**, Expression t-SNE maps of hepatocyte marker genes *CLU* and *SERPINA1*. **c**, Expression t-SNE maps of *LYZ*, *PIGR*, *MUC5A*, *AGR2*, *AKR1B10*, *CCND1*, *H25AZ*, and *HIST1H1A*.

**Extended Data Figure 9.**
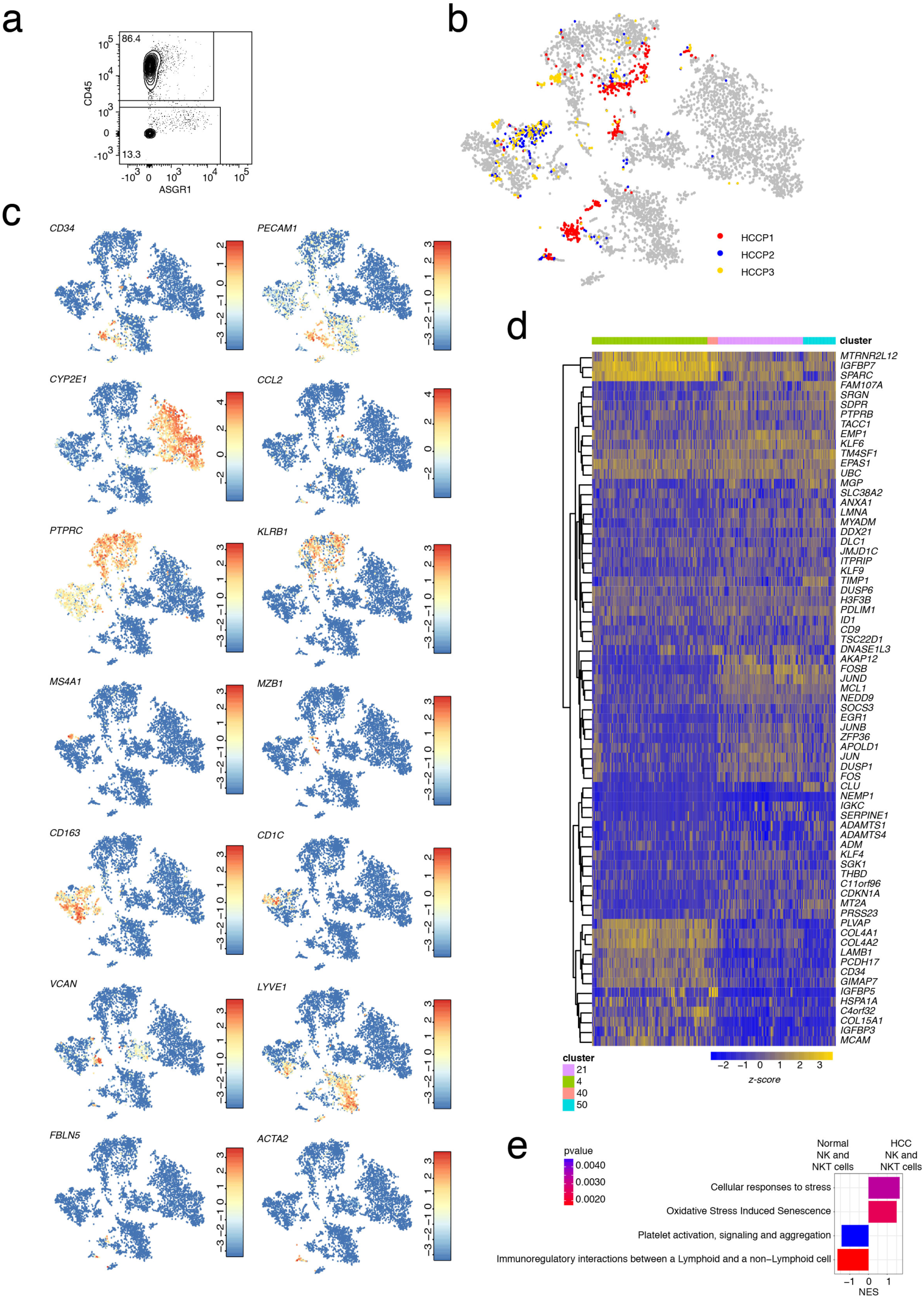
Cell types from patient-derived HCC exhibit perturbed gene expression signatures. **a**, FACS plot of CD45 and ASGR1 staining on cells from HCC samples. **b**, Symbol t-SNE map showing the IDs of the HCC patients. **c**, Expression t-SNE maps of *CD34*, *PECAM1*, *CYP2E1*, *CCL2*, *PTPRC*, *KLRB1*, *MS4A1*, *MZB1*, *VCAN*, *LYVE1*, *FBLN5*, *ACTA2*. **d**, Heatmap of differentially expressed genes between normal *CD34*^+^ MaVEC clusters and HCC endothelial cells from HCC clusters (Benjamini-Hochberg corrected *P*<0.05). **e**, GSEA for differentially expressed genes between normal and HCC-resident NK and NKT cells (Benjamini-Hochberg corrected *P*<0.05). The bar chart shows the normalized enrichment score (NES) and highlights the p-value.

**Extended Data Figure 10.**
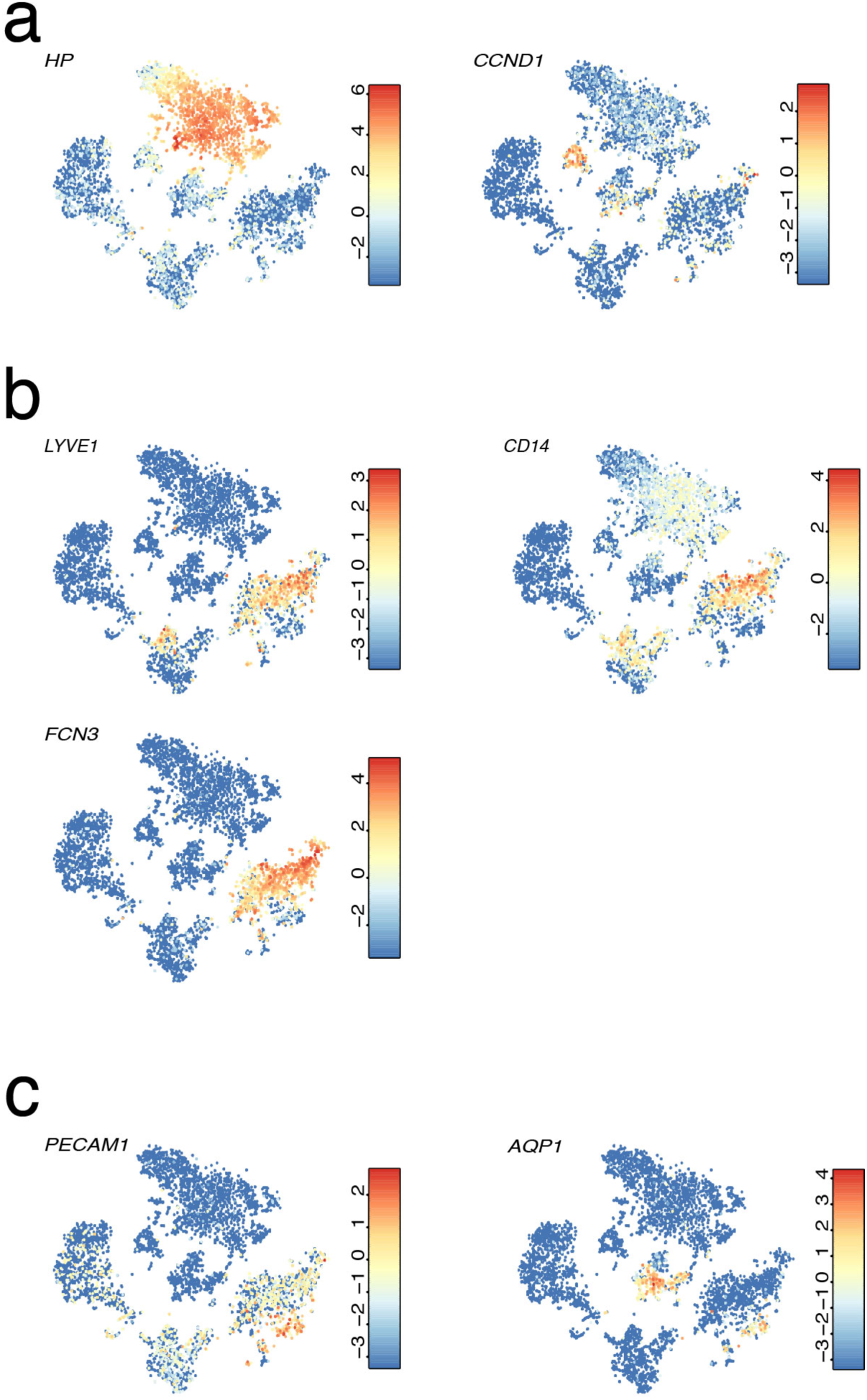
Transplanted human liver cells in a humanized mouse model exhibit a distinct gene signature compared to cells within the human liver. **a**, Expression t-SNE maps of *HP* and *CCND1*. **b**, Expression t-SNE maps of the LSEC zonated genes *LYVE1*, *CD14* and *FCN3*. **c**, Expression t-SNE maps of *PECAM1* and *AQP1*.

